# Metabarcoding a Metacommunity: detecting change in a wetland wilderness

**DOI:** 10.1101/819714

**Authors:** A. Bush, W.A. Monk, Z.G. Compson, D.L. Peters, T.M. Porter, S. Shokralla, M.T.G. Wright, M. Hajibabaei, D.J. Baird

## Abstract

The complexity and natural variability of ecosystems present a challenge for reliable detection of change due to anthropogenic influences. This issue is exacerbated by necessary trade-offs that reduce the quality and resolution of survey data for assessments at large-scales. The Peace-Athabasca Delta (PAD) is a large inland wetland complex in northern Alberta, Canada. Despite its geographic isolation, the PAD is threatened by encroachment of oil sands mining in the Athabasca watershed, and hydroelectric dams in the Peace watershed. Methods capable of reliably detecting changes in ecosystem health are needed to evaluate and manage risks. Between 2011 and 2016, aquatic macroinvertebrates were sampled across a gradient of wetland flood frequency, applying both microscope-based morphological identification, and DNA metabarcoding. Using multi-species occupancy models, we demonstrate that DNA metabarcoding detected a much broader range of taxa and more taxa per sample compared to traditional morphological identification, and was essential to identifying significant responses to flood and thermal regimes. We show that family-level occupancy masks high variation among genera, and for the first time, quantify the bias of barcoding primers on the probability of detection in a natural community. Interestingly, patterns of community assembly were near random, suggesting a strong role of stochasticity in the dynamics of the metacommunity. This variability seriously compromises effective monitoring at local scales, but also reflects resilience to hydrological and thermal variability. Nevertheless, simulations showed the greater efficiency of metabarcoding, particularly at a finer taxonomic resolution, provided the statistical power needed to detect change at the landscape scale.

## Introduction

Tackling the global loss of biodiversity (1) is hindered by a lack of basic biological information needed to guide sustainable management strategies (2). Despite legal protections, freshwater ecosystems are increasingly degraded by multiple stressors (3). In addition, the quality and volume of data collected by monitoring programs often fails to support evidence-based management decisions (4–6). Here, we demonstrate how DNA metabarcoding can resolve challenges faced by traditional monitoring, alter our perspectives on ecosystem dynamics, and improve our understanding of natural variation and sampling error, supporting evidence-based decision making.

DNA barcoding uses short genetic sequences to identify individual taxa. By contrast, DNA metabarcoding supports simultaneous identification of entire assemblages via high-throughput sequencing (7, 8). Using metabarcoding for ecosystem monitoring provides an opportunity to identify organisms in bulk samples at a high taxonomic resolution consistently and accurately (“Biomonitoring 2.0”; 9). The accuracy, consistency, and resolution of taxonomic identification remains a constraint for many biomonitoring programs that must trade-off data quality to make assessment protocols rapid and cost-effective (10). Aquatic macroinvertebrates exemplify this challenge, as their diversity of forms and functions are sensitive to multiple drivers of ecosystem condition. Thus, ecosystem degradation can be identified based on changes in assemblage composition due to environmental filtering (5). Despite decades of development, the challenges associated with traditional methods of sample processing limit inference of biomonitoring programs to gross status classifications (e.g. 11). Metabarcoding presents an opportunity to describe community composition more accurately and consistently, supporting more effective and informative biomonitoring (12, 13).

The Peace-Athabasca Delta (PAD) in northern Alberta, Canada (Figure 1; and 14). The PAD is North America’s largest inland delta (approximately 6,000 km^2^), and is located at the confluence of the Peace and Athabasca Rivers, consisting of hundreds of lakes and wetlands that become connected during flood events, particularly when spring snowmelt leads to ice-jams (15, 16). The PAD is a Ramsar wetland, protected within Wood Buffalo National Park, a UNESCO World Heritage site. Nonetheless, there have been concerns that the PAD could be affected by upstream developments, including current and proposed hydroelectric dams on the Peace River, continued expansion of oil sands mining on the Athabasca River to within 30km of the park boundary, and climate change (17). Assessing how such factors influence the integrity of a natural wilderness is made more challenging by the paucity of biological surveys that have been conducted, and the logistics of working in such a remote region. To gain a better understanding of the PAD’s ecology, rapid assessments of aquatic macroinvertebrates have been conducted in since 2011 to establish a baseline understanding of the ecosystem’s diversity and structure (14, 18). Importantly, while surveys have followed established protocols from the Canadian Aquatic Biomonitoring Network (hereafter CABIN)(19), samples were processed using both traditional and DNA metabarcoding approaches, allowing us to test the power of each approach to support environmental management of the delta for the first time.

**Figure 1.**
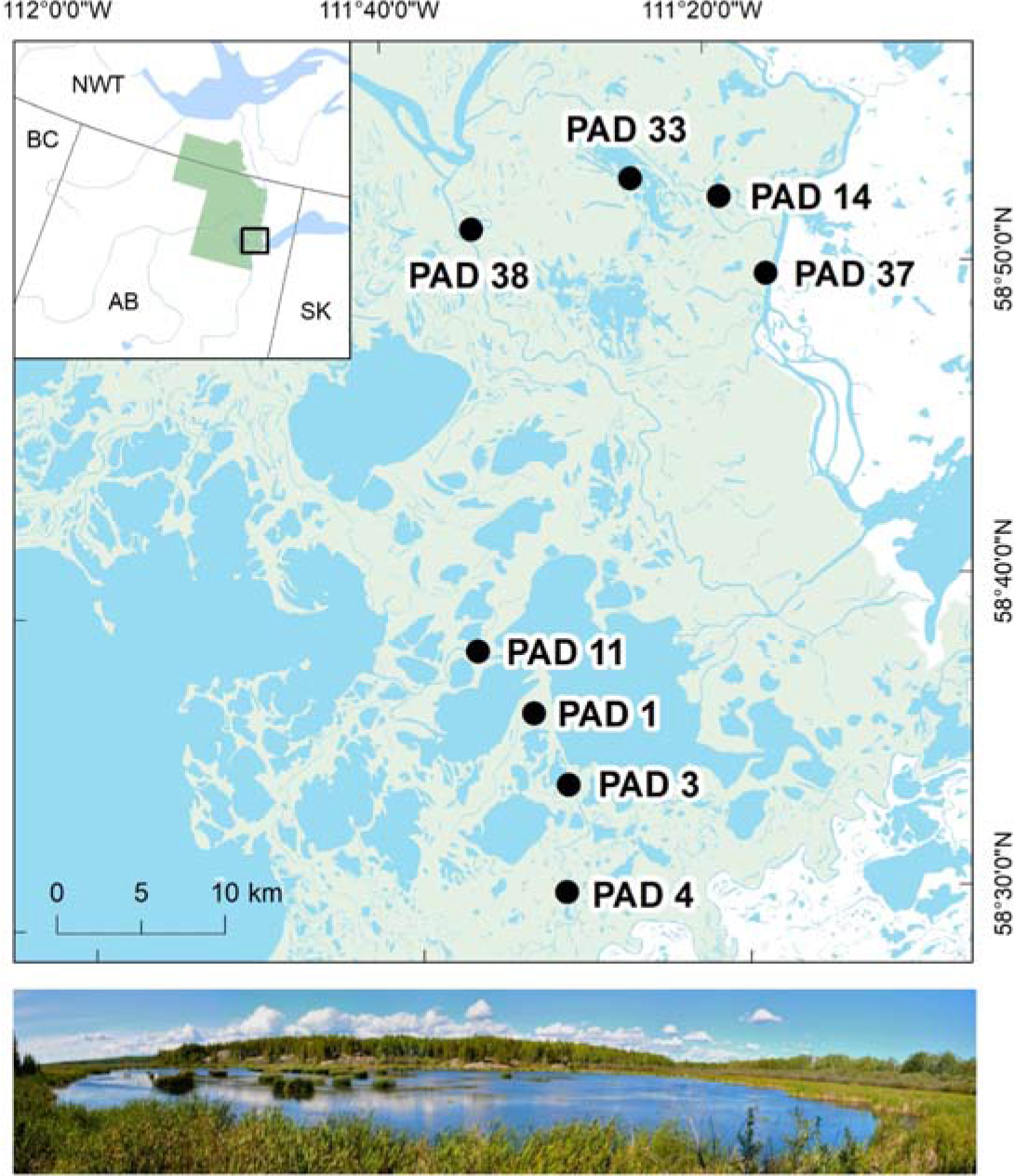
Figure 1: Location of sampling sites in the Peace-Athabasca Delta. The inset shows the full extent of Wood Buffalo National Park in Alberta (AB), and boundaries of neighbouring provinces: British Columbia (BC), Saskatoon (SK), and the Northwest Territories (NWT). Photo taken at Rocher River wetland (PAD 37).

Sampling error is a ubiquitous feature of any ecological survey, irrespective of the methodology, and of particular concern is the frequency of false absences (20). Depending on the covariance of species’ detectability with other environmental characteristics, models can be structurally biased and their confidence overestimated (21). Although imperfect detection is very common, and freshwater biomonitoring protocols have a long history of standardization (5), there are few examples of research explicitly quantifying the nature of sampling error (e.g. 22), instead including it within natural sources of variability (i.e. as “noise”; 23). An alternative is to specify the likelihood of detection (the observation process model) and simultaneously correct our estimates of species occurrence (the ecological state model) within a single hierarchical framework (24). In this study, we employed multi-species occupancy models (MSOMs; 25, 26) to account for the effects of imperfect detection on estimates of macroinvertebrate diversity, drawing upon data from six years of macroinvertebrate surveys in the PAD. We quantify the efficiency with which the macroinvertebrate community can be surveyed using both traditional morphological identification and DNA metabarcoding, and demonstrate that these approaches make a qualitative difference to our view of the metacommunity is structured, to the efficiency of monitoring, and consequently our power to detect change (27).

## Results

A key difference between our sampling approaches was that the standard CABIN wetland protocol (19) provided estimates of relative abundance based on counts from a subset of each sample, whereas sequences identified using DNA metabarcoding were converted to presence-absence data (13, 28). In addition, CABIN identified 74 families based on morphological features, but metabarcoding could identify 109 families, as well as 263 genera (see Supplement 1 and Fig. S1.6). As a result, we trained four hierarchical MSOMs for each data type: 1) counts of macroinvertebrate families from CABIN (CABIN *Fcount*), 2) the presence-absence of macroinvertebrate families from CABIN (CABIN *Fpa*), 3) the presence-absence of macroinvertebrate families from DNA data (DNA *Fpa*), and 4) the presence-absence of macroinvertebrate genera from DNA data (DNA *Gpa*). Although metabarcoding can discriminate among taxa at even finer resolution (i.e. species), given the prevalence was lower than the prevalence of genera and the available sample size, we did not feel the detectability and occupancy of those taxa could be estimated reliably.

### Occupancy and detectability

The CABIN *Fcount* model predicted total abundance was dominated by four taxa (two Chironomidae subfamilies, Oligochaeta and Planorbidae), but also suggested that almost all taxa were present everywhere within the PAD (i.e. site occupancy ≈ 1), with no environmental covariates retained in the final model. This scenario is plausible, but if we apply the predicted probabilities of detection and same survey effort (number of individuals counted), and assume taxa are sampled at random from the pool of individuals, the CABIN *Fcount* model suggested we should have observed 38 taxa on average instead of 18. Non-random aggregation of individuals is typical of ecological communities (29), and may be why the model appeared to be misspecified.

In contrast to the count-based model, the presence-absence models all suggested taxon site-occupancy was below 1 (Fig. 2, Fig. S1.10), although the “U-shaped” form of the hyperparameters in Fig. 2a and 2c was an artefact of the bounded distribution (29). The CABIN *Fpa* model estimated the probability of detecting macroinvertebrate families was lower than the DNA *Fpa* model (Fig. 2b right-skewed relative to 2d,e; see also Figure 3). Models must balance the expected occupancy to fit with the detections, and probability of detection made in each survey, and the CABIN *Fpa* model therefore also predicted higher occupancy than the DNA *Fpa* model (Fig. 2a left-skewed relative to 2c). The differences in occupancy and detectability of specific families were not associated with prevalence, although many taxa were not recorded by both approaches, and therefore cannot be compared (red points in Fig. 3; see Supplement 1 for detail). In addition, detectability using DNA metabarcoding is intrinsically linked to the genetic primer used (30), and the importance of primer bias is well known from mock laboratory samples (e.g. 31). Here we show biases in detectability can be quantified as part of the observation model, either at the community-level (Fig. 2d and 2e), or for individual taxa (Fig. S1.11). Lastly, neither the CABIN *Fcount* or *Fpa* model retained environmental variables to explain changes in occurrence, whereas both DNA occupancy models did so consistently. The covariates retained were 1) the frequency of spring and summer floods (i.e. connections between the wetland and river), 2) time since the ice melt, and 3) maximum water temperature prior to each survey. Responses to environment at the community-level were almost neutral (Fig. S1.13), and the posterior distribution of coefficients differed from zero for only a minority of taxa (Fig. S1.14), but their inclusion in the model suggests the high inter-annual turnover (Fig. S1.7) may be explained in part by deterministic factors.

**Figure 2.**
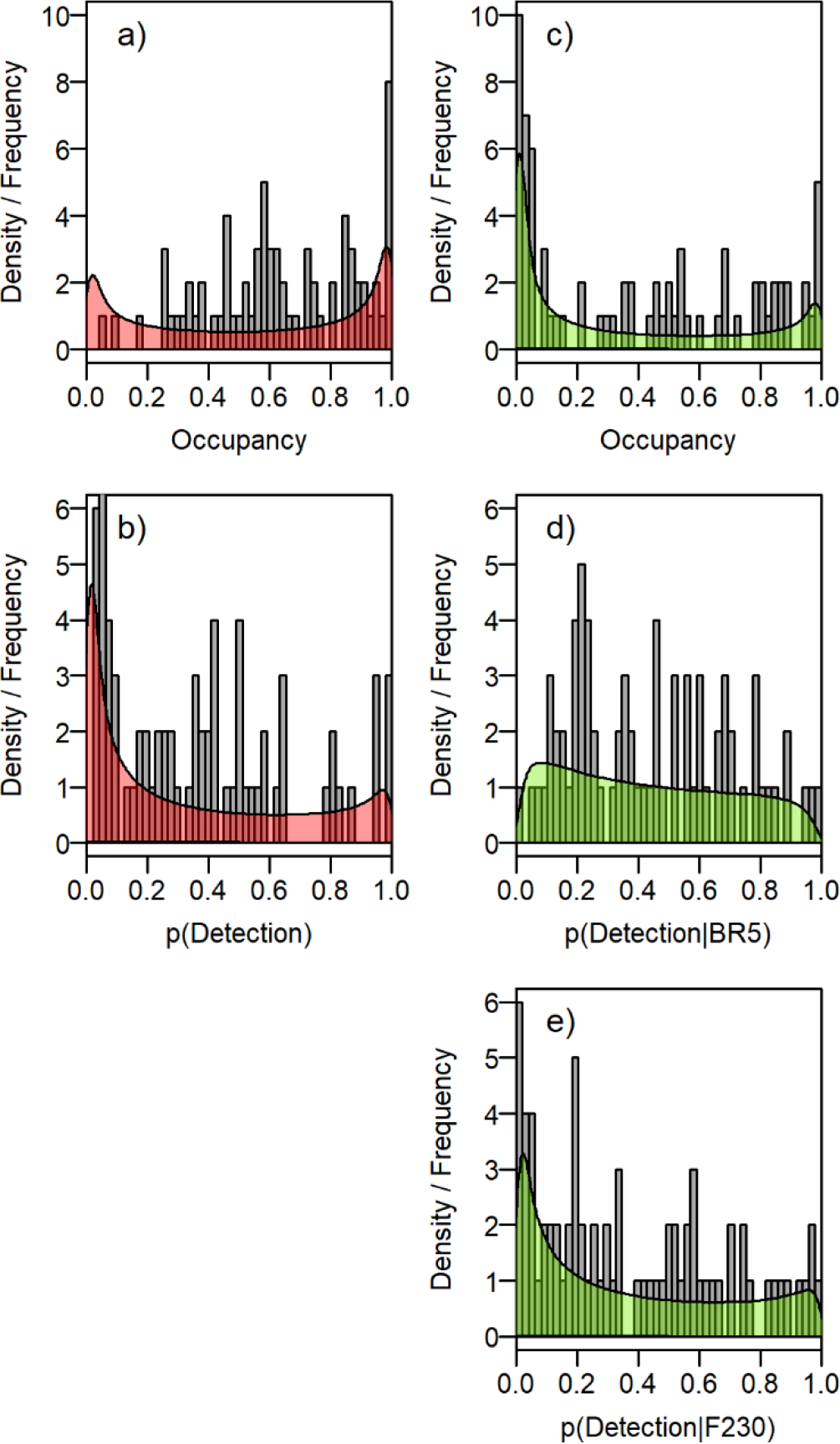
Predicted occupancy (a,c) and detectability (b,d,e) of taxa based on the presence-absence data collected using the CABIN protocol (a,b) and DNA metabarcoding (c-e) at the family-level. Detectability using metabarcoding is further split by primer pair (d,e). The shaded polygons describe the probability density of the community hyperparameters, and the grey bars indicate the underlying frequency of the values estimated for each taxon. See Figure S.10 for the CABIN *Fcount* and DNA *Gpa* model distributions.

**Figure 3.**
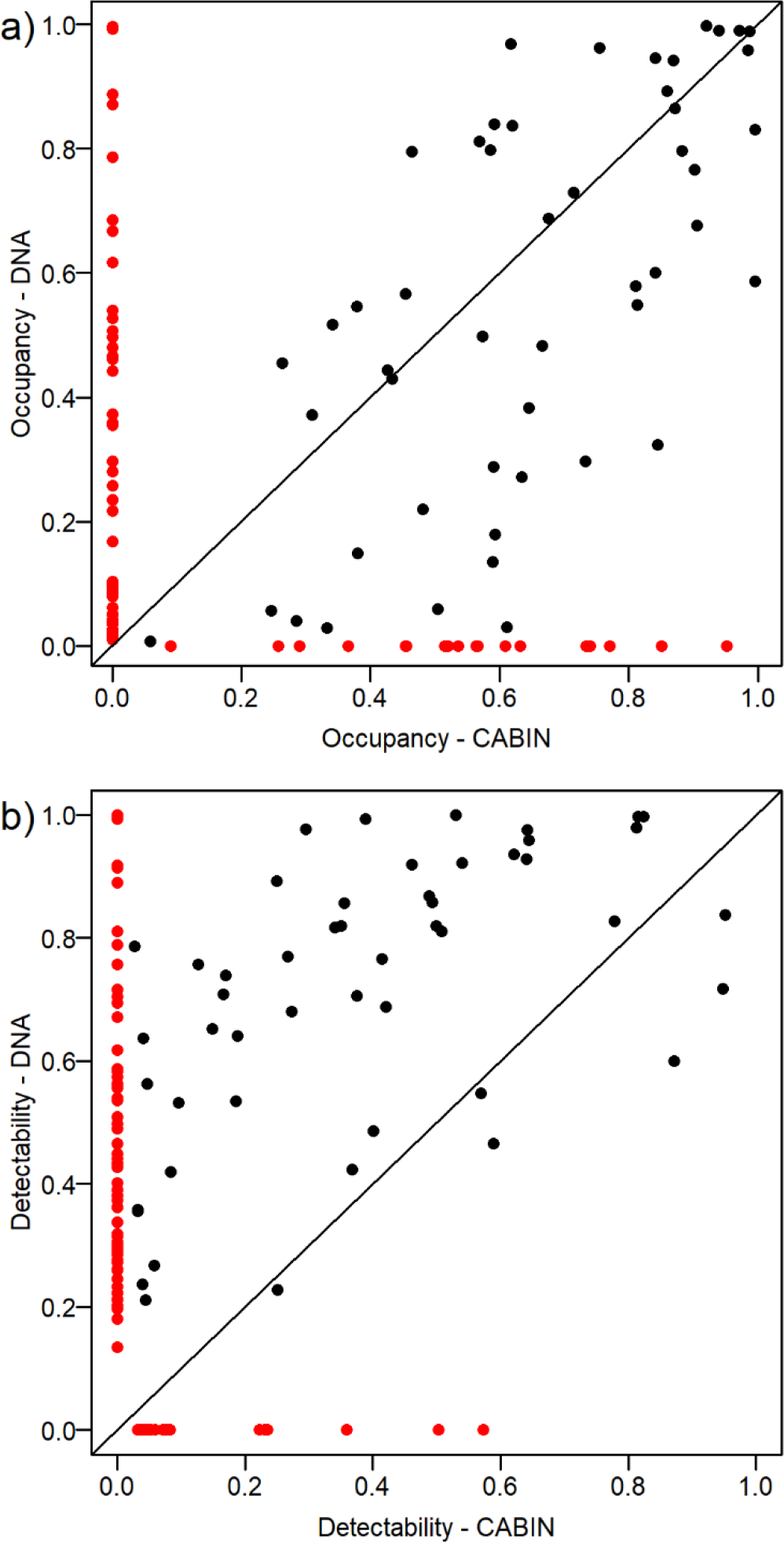
Comparison of a) occupancy and b) detectability estimates in models trained by CABIN data and DNA metabarcode data at the family-level (n= 50). Red points indicate taxa not observed by the complementary method i.e. 18 and 59 families were unique to CABIN and metabarcoding respectively. See Supplement 1 for further information on the identities of unique taxa.

### Alpha, beta and gamma diversity

Recognizing that imperfect detection is commonplace in ecological surveys, it follows that regional (gamma diversity, Fig. S1.8) and local (alpha diversity, Fig. S1.9) diversity is routinely underestimated. As the CABIN *Fcount* model effectively assumed alpha and gamma diversity were equal, it estimated that only two families were likely to have gone undetected in the metacommunity. Conversely, the CABIN *Fpa* model estimated ~20 families were missed (i.e. γ = 95), a 28% increase on the observed total. Interestingly, this estimate was still short of the richness observed using metabarcoding (n=109), and based on the distribution of detection probabilities, the DNA *Fpa* and *Gpa* models estimated the metacommunity could potentially contain 130 families and 360 genera, a 19% and 37% increase (Fig. S1.8).

Although imperfect detection always underestimates richness, its effect on the observed compositional dissimilarity between sites (beta diversity) is less predictable. The observed pairwise-dissimilarity of samples consistently exceeded 40%, both within and between years, with no consistent increase over time (Fig. S1.7). Our analysis showed that compositional turnover in the CABIN dataset was overestimated, whereas for the DNA models the corrected and observed dissimilarities were similar (Fig S1.15), although temporal turnover (i.e. inter-annual, within-site dissimilarity) was marginally overestimated by the DNA dataset. This implies that although metabarcoding underestimated alpha diversity at each site, the proportions of the taxa missed that were shared or unique to site-pairs were similar. Finally, one predictable aspect of turnover is that as the taxonomic resolution is increased, sub-taxa are on average less prevalent than their parental ranks (see Fig. S1.12a), typically harder to detect (presumably because they are also less abundant than parental ranks), and therefore dissimilarity among sites at the genus-level was 7% higher compared to the family-level.

### Power analysis

The power to detect statistically significant changes depends on the strength of the ecological signal relative to other natural variability, as well as the efficiency with which we can accurately describe ecological state, factors directly related to taxonomic resolution, and detectability (27). We simulated the PAD metacommunity based on a fitted distribution of occupancy and estimated gamma diversity to represent its baseline condition, and then took subsamples that reflected the observed biases in each sampling approach. Note that the true state and behaviour of the system are unknown, and underlying processes were instead inferred by the MSOMs after quantifying observation biases. Human impacts that might affect the PAD system in the future were also unknown, and this analysis therefore aimed to identify our power to observe a generalized stressor effect. To keep the process model consistent, we based simulations on the most detailed DNA *Gpa* model, and then aggregated taxa to higher ranks to compare power among sampling approaches. A complete description of the simulation and power analysis is provided in Supplement 2.

A natural consequence of high, near-random, background variation in composition is that degradation of a wetland site would need to be severe to raise concerns. Instead, it is more effective to measure when there is a shift away from our expectation of the PAD metacommunity aggregated across sites (i.e. changes in occupancy of many taxa). Even so, based on the high natural variability of the PAD, the survey effort needed to confidently detect shifts in occupancy in any year would be prohibitive. As a result, we considered a monitoring system to be adequate if significant differences in composition were detected within 2 years (at least 50% of the time, Fig. S2.4). Our results demonstrated that our power increased: 1) as the number of sites sampled increased (but the rate of increase declined beyond 8-10 sites per year); 2) with DNA metabarcoding compared to CABIN sampling, and with genus- compared to family-level data; and 3) if we sampled sites multiple times (but gains depended on the number of sites and sampling approach) (Fig. 4). Statistical power also varied by stressor type because metacommunity shifts were readily apparent if the stressor impacted prevalent taxa, whereas changes were challenging to observe if prevalent taxa were also tolerant. The relationship between taxon occupancy and their sensitivity to a stressor was therefore most influential when sample sizes, and hence our power to detect rare taxa, were low (Fig. S2.6).

**Figure 4.**
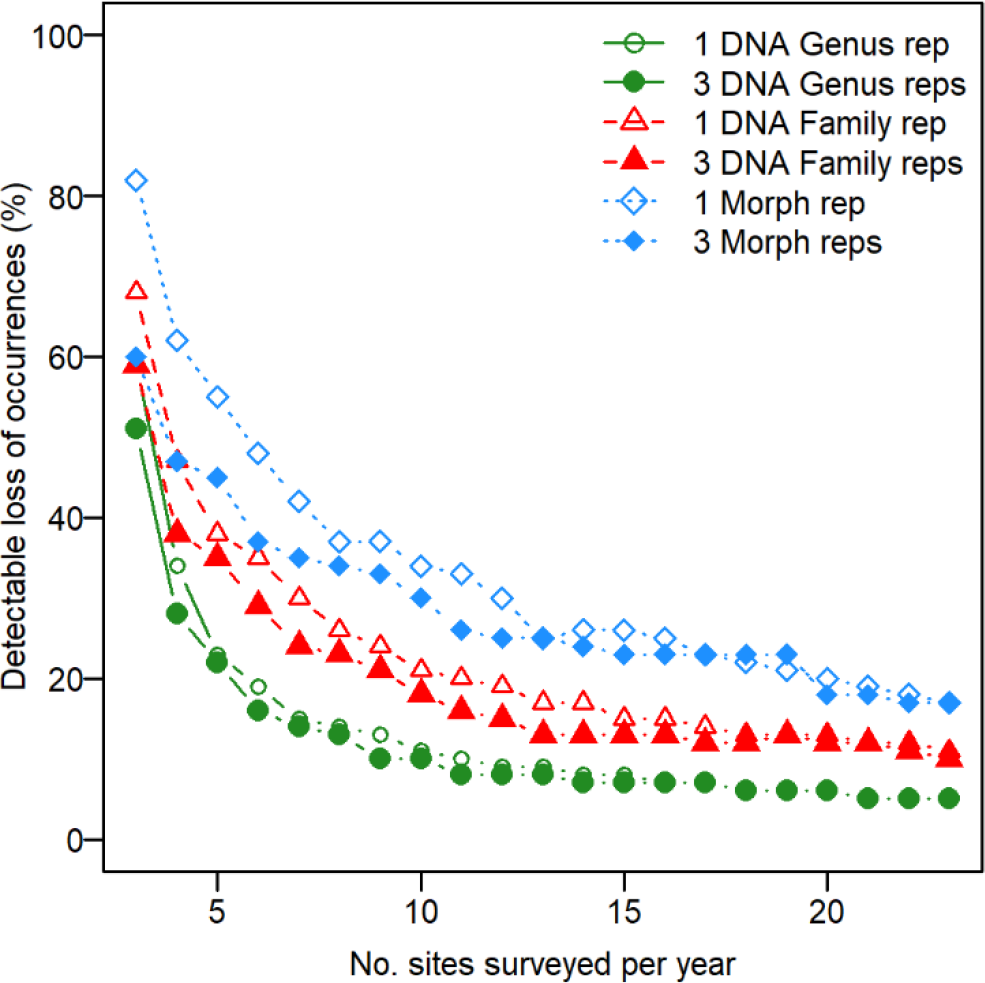
Minimum reduction to community occupancy that is detectable >50% of the time with 95% confidence in response to number of sites surveyed annually. Lines show the average of 100 simulations based on the CABIN *Fpa* (blue), DNA *Fpa* (red) and DNA *Gpa* (green) occupancy-detection models, with either single (open symbol) or triplicate (closed symbol) samples per site. Taxon tolerance was not correlated with occupancy. See Supplement 2 for further information.

## Discussion

The Peace-Athabasca Delta represents one of Canada’s national biodiversity treasures. Yet multiple external pressures, including the development of oil sands, hydroelectric power, wildfires and climate change are potentially affecting biodiversity through modification of natural physical processes in the area and threaten its World Heritage listing (17). Our study demonstrates that the PAD is an immensely rich habitat, including over 25% and 20% of all aquatic macroinvertebrate families and genera recorded by the CABIN national biomonitoring program (32). Furthermore, communities exhibit near-random patterns of spatial and temporal turnover, a property rarely observed in freshwater systems (33). As a result, impacts on the wetland macroinvertebrate community are difficult to establish at local scales because occurrence is weakly related to environmental factors and site composition can fluctuate rapidly over time (Fig. S1.7). Properties of the metacommunity must therefore be aggregated across sites, and directional shifts can only be inferred when dissimilarities are unlikely to be explained by stochastic differences in our null baseline model (34). Our analysis shows that detecting a decline in metacommunity condition would depend on both sample size and stressor type, and that further changes to sampling design may be required to detect change earlier or at specific locations of concern.

The most significant finding of this study was the value added to biomonitoring data generated by DNA metabarcoding of bulk community samples. Our analysis supports previous studies that have shown the breadth and resolution of taxonomic information achievable with metabarcoding (e.g. 14, 35). Clear differences in occupancy and detectability profiles with metabarcoding (Fig. 3) influence our description of baseline reference conditions (36, 37). Further differences in estimates of occupancy with increasing taxonomic resolution (Fig. S1.12) may also indicate differential environmental responses (38, 39). We did not find evidence to suggest the count data (’relative abundance') in CABIN samples were necessary to detect changes in ecological structure. In fact, only the presence-absence DNA metabarcoding models identified significant relationships with the major environmental covariates of this region (16). These effects could be estimated precisely because detectability, and thereby sampling efficiency, was higher for so many taxa using DNA metabarcoding (13). We also used the occupancy model framework to compare detectability of each taxon with different primers, a more robust measure of their complementarity than lists of taxa observed. Quantifying detectability is vital to making the results of this study comparable to others with varying protocols, and this approach could be used to refine and select complementary primers (28, 31). Crucially, DNA metabarcoding, particularly at genus-level, substantially improved our power to detect ecosystem-scale changes compared to traditional CABIN sampling (Fig. 4, Supplement 2).

A second outcome was the importance of explicitly considering imperfect detection. Practitioners are well aware of sampling differences (e.g. 40), but have typically focused on how those errors propagated to aggregated metrics, rather than explicitly quantifying the sources of uncertainty (23). Hierarchical occupancy models accommodate irregularly sampled data, estimate community properties (that extend inference to rare taxa), and allow straightforward biological interpretations of those parameter estimates (41). There have been few examples of hierarchical models accounting for detectability in freshwater ecology (e.g. 42, 43), despite studies showing it can bias our interpretation of taxonomic, functional, and phylogenetic diversity at the community-level (e.g. 44). Given the high prevalence of false absences it is not surprising occupancy models are becoming commonplace for analysing eDNA data (45), although it appears multi-species models are still rare (46). Importantly, what these and other studies have shown is that the time and expense of adding replicate samples may be the most efficient way to improve the statistical power of a study (24, 47). Inferences about the number of taxa missed in the metacommunity naturally carry some uncertainty (48), but by acknowledging imperfect detection, risks can be quantified and decision makers’ overall efficiency can be improved. This analysis minimises the likelihood of management agencies responding to a false signal of degradation (Type 1 error), and identifies how to optimize survey design to ensure we have the necessary power to detect a desired degree of change (Type 2 error) (27, 49).

Although our analysis provided evidence of environmental filtering, the distribution of beta diversity was equivalent to that expected from random assembly, suggesting that the metacommunity was operating in a quasi-neutral manner at the scale of our analysis (50). Quasi-neutral dynamics are expected to be commonly observed in taxon-rich communities, but given the high degree of landscape connectivity, we would expect mass effects, rather than dispersal limitation, to underlie the low habitat specificity of the community (51). Models of coexistence that combine stochasticity with niche theory may therefore better explain the structure and dynamics of aquatic invertebrate communities in the PAD without relying on the fragile premise of ecological equivalence in neutral theory (50, 52). Although many ecologists have acknowledged stochastic processes are likely to have a role in understanding community composition (53, 54), we are unaware of any biomonitoring programs that incorporate, or even acknowledge community assembly mechanisms other than environmental filtering (e.g. 37). Our results firmly challenge that traditional perspective, and if we wish to understand the resilience of the PAD, we must adopt a metacommunity perspective (55). More broadly, a metacommunity perspective of the PAD could indicate which assembly processes are absent from more managed landscapes, therefore providing critical insights into the mitigation of biodiversity loss at the landscape scale.

The isolation of wilderness areas like the Peace-Athabasca Delta implies a pristine nature, but that isolation has also hindered our appreciation of the sheer magnitude of diversity which occurs there, and has until now precluded a basic description of how community structure changes over space and time. Near-random patterns of assembly and substantial sampling error pose a challenge to detecting ecosystem change. Without evaluating data quality and statistical power at the start, many monitoring programmes are unable to confidently reject a false null hypothesis, undermining project goals and providing a misleading sense of achievement (27, 56). Despite the high turnover, we show the statistical power of data generated by DNA metabarcoding was superior to traditional biomonitoring approaches for the detection of large-scale ecosystem change. Although macroinvertebrate composition provides a wealth of information, the power to detect and draw inference from taxonomic changes will be improved by further refining the list of taxa that respond to particular threats (e.g. oil sands contaminants), particularly by linking metabarcoding to trait databases (57), and this remains a major focus of our ongoing research.

## Materials and Methods

### Field surveys

Field survey methods followed the CABIN wetland macroinvertebrate protocol (14, 19). Briefly, aquatic invertebrates were sampled by sweeping submerged and emergent aquatic vegetation at wetland edges for 2 minutes. A sterile 400μm mesh net was steadily moved in a zig-zag pattern, from the surface of the sediment to the water surface, to capture disturbed organisms and minimise the amount of sediment collected. Excess vegetation was carefully rinsed and removed, and samples with excess sediment were sieved. Material was placed in sterile 1L polyethylene sample jars, filled no more than half full, and immediately preserved in 95% ethanol in the field. Samples were stored in a cooler with ice in the field and transferred to a freezer at the field station before shipment. Nets were disinfected between each new site, and field crews wore nitrile gloves to collect and handle samples, minimizing the risk of cross-site contamination.

### Sample Processing

In total, 126 and 138 samples were collected from 72 separate site visits for the CABIN and DNA metabarcoding datasets, respectively (Supplement 1, Table S1.2). Samples identified using morphological characteristics were processed and identified in accordance with the CABIN laboratory manual (19). Briefly, material from each 2-minute sweep was subsampled using a 100-cell Marchant box. Successive cells were processed until at least 300 individuals were identified and a minimum of five cells were processed. Most taxa were identified to family level, although for some groups only class- or order-level identification was recovered, and given the importance and diversity of Chironomidae, we retained four sub-family divisions that could be reliably identified (Supplement 1).

The lab protocol for processing samples for DNA metabarcoding followed the same procedure as outlined in Gibson, et al. (14). This targeted the CO1 amplicon using two complementary primers, BR5 and F230R (310bp and 230bp respectively; 30, 58). All DNA samples were analysed using BR5, although F230R was only introduced after 2011. While field and lab protocols have remained consistent since the study began, there have been a number of advances made in bioinformatic tools, as well as expansion of the reference sequence libraries supporting the identification of taxa (59). The bioinformatic pipeline used to process all samples in this study, as well as the CO1 classifier that allocates sequences to the most likely taxa, is described in Supplement 1 and available on Github (60).

### Hierarchical MSOM

Multi-species occupancy models (MSOMs) employ a flexible hierarchical framework that allows for imperfect detection to predict species’ occurrence (25). Our analysis adapted the notation and code provided by (41) as the basis for this study (see Supplement 1):.

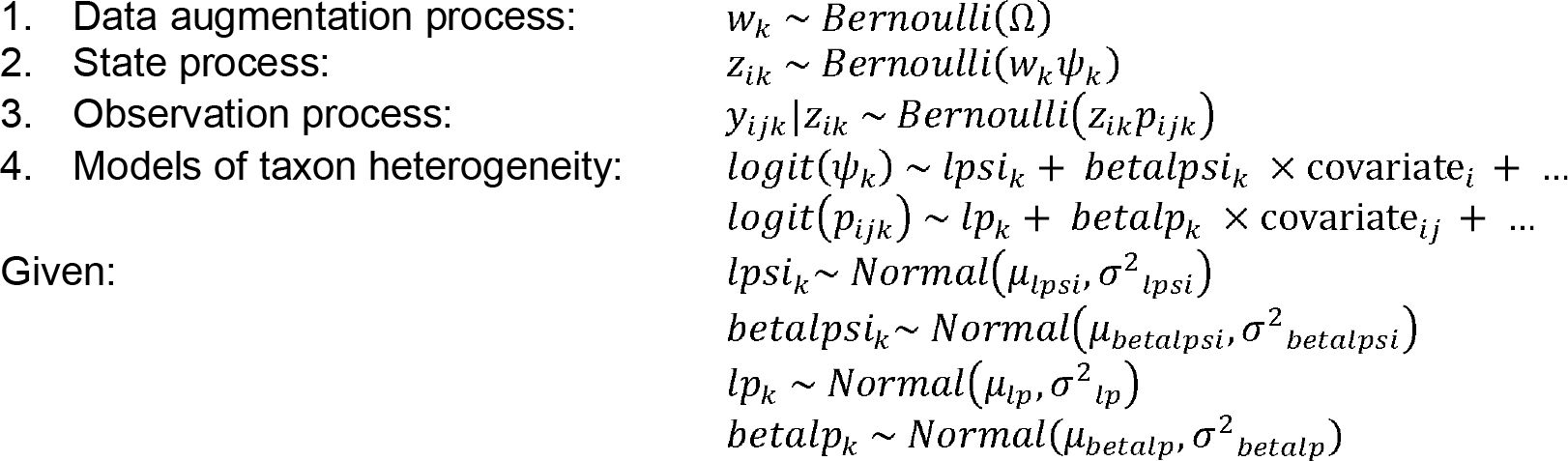

The observed data yijk describe the detection or non-detection of taxon k at site i in replicate sample j. Replicate observations, in our case simultaneous independent samples (21), allowed the model to discriminate between processes that determine the system’s state (occupancy) and the observation process (detectability). The occupancy of each taxon at each site zik is described by a Bernoulli trial with probability Ψik, and the likelihood of detecting the respective taxa in each replicate sample is described by another set of Bernoulli processes with probability pijk. Changes in occupancy and detectability due to various covariates were also tested using multiple logistic regression. Individual intercepts and slopes represented species-specific random-effects, governed by a common prior distribution whose mean and variance were estimated as a community-level hyperparameter.

The statistical distributions of the parameters governing occupancy and detectability shared by the community were used to consider the possibility of other taxa in the metacommunity that were not recorded in any visit to any site, a process known as data augmentation (48, 61). Given a sufficiently large total pool of M potential taxa, a set of binary indicators wk, governed by the parameter Ω, represent the probability each taxon is part of the community. The total number of taxa in the metacommunity (γ-diversity) is therefore simply the sum of wk.

The occupancy model above was suitable for presence/absence observations of taxa, but CABIN samples also included information on the relative abundances of taxa. To utilize all information available, we constructed a community-level N-mixture model that estimates the latent abundance Nik of each taxon, rather than their occurrence (zik), and modelled counts as a function of a Poisson distribution.

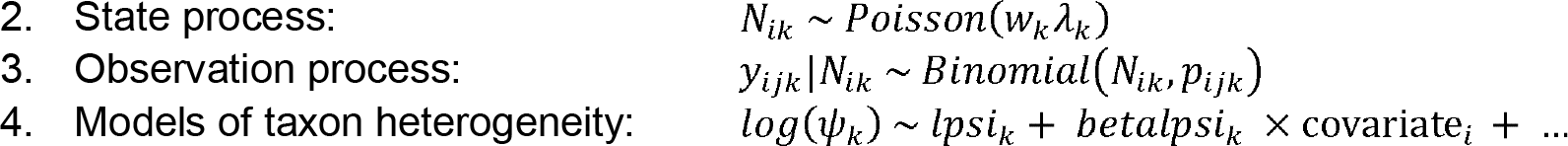

Finally, model selection for covariates of taxon heterogeneity in both the occupancy and N-mixture models was determined by a set of binary indicator variables Vx1-xn, one for each of the n predictor variables used (62). Using Vx ~ Bernoulli(0.5) as standard priors, variables had an equal likelihood of being included or excluded from likelihood estimates, and model selection was therefore based on which combination had the highest joint posterior probability p(Vx1-xn=1). Note convergence of the Vx indicators was very slow, particularly in the most complex models, and a “slab and spike” approach did not improve mixing (see 7.6.2 in 41).

Analyses were conducted using the R package jagsUI (63). We assessed model convergence of all monitored parameters across chains by visual inspection of trace plots and by using the Gelman-Rubin statistic (64), with the diagnostic value <1.1. As over-dispersion cannot be estimated from the binary responses in occupancy models (41), plots of Dunn-Smyth residuals for fitted estimates of occupancy and detectability were used to evaluate the fit of separate taxa (65). Although plots suggest the models were well fit in most cases, the pattern of residuals suggested there may have been other covariates, or non-linear effects, missing from the models influencing the occupancy of some taxa.

### Simulation and power analysis

A hypothetical presence-absence matrix of the metacommunity was derived from estimates of gamma diversity and occupancy in the DNA *Gpa* occupancy model from which we could manipulate sampling designs. Environmental covariates were varied according to the mean and standard deviation of values observed from the surveys available to us (Figure S2.1), but the simulation was not spatially explicit. Whilst occupancy covariates drove some temporal turnover (Figure S2.2, approximately 10-27%), this was insufficient to replicate the turnover observed (Fig. S2.3), so we used permutations of the presence-absence matrix to simulate further stochastic changes in composition (i.e. local extinction/colonization; 66). Replicating observed turnover required the complete redistribution of occurrences (i.e. random assembly patterns). Taxon occupancy (row sums) and site richness (column sums) were held constant during permutation. The metacommunity was modified by successively removing occurrences of taxa based on a hypothetical distribution of tolerances, which were themselves generated to covary with the distribution of occupancy. Sampling error was applied by a binomial function weighted by the taxon’s probability of detection, and the ‘detected’ composition of reference and modified metacommunities were then compared using mvabund (67). Power of DNA *Gpa* was compared to DNA *Fpa* and CABIN *Fpa* approaches by aggregating genera to family-level, and subsequently applying the family-level detection probabilities. See Supplement S2 for more details.

## Acknowledgments

We acknowledge support for this work from a Large-Scale Applied Research Project (LSARP) award from Genome Canada to MH and DJB. DJB also received support from the NSERC Discovery Grants Program, and through Environment and Climate Change Canada program funds, including the Genomics Research & Development Initiative that supported both AB and TM. Access to sites was supported by the Canada-Alberta Oil Sands Monitoring Program. This work was funded in part under the Oil Sands Monitoring Program, and is a contribution to the Program, but does not necessarily reflect the position of the Program. Field support was provided by Parks Canada (Jeff Shatford, Queenie Gray, Jason Straka, Sharon Irwin and air-boat pilots Ronnie and David Campbell) and Environment and Climate Change Canada’s Watershed Hydrology and Ecology Research Division (Kristie Heard, Colin Curry, Daryl Halliwell, Tom Carter, Adam Bliss, Adam Martens, Cath Choung and Steff Connor).

