## Supplement 1 for "Metabarcoding a Metacommunity: detecting change in a wetland wilderness"

### Supplementary Material S1: Multi-species occupancy modelling of freshwater macroinvertebrates in the Peace-Athabasca Delta based on morphological identification and DNA metabarcoding.

*Alex Bush*

*26th June 2019*

#### Contents


**Input Data** **2**
- Sampling sites 2
- Environmental parameters 2
  - Temperature 3
  - Flood frequency and duration 4
- Assemblage Data 6
  - Morphological Data 6
  - DNA Metabarcoding Data 6
  - Exploration of community data 8

**Hierarchical Model Framework** **10**
- Occupancy Modelling 10
  - JAGS Models 10
  - Convergence, model fit and variable selection 13
  - N-mixture Model 14

**Results** **16**
- Metacommunity size (gamma diversity) 16
- Site richness (alpha diversity) 17
- Occupancy and detectability 18
  - CABIN vs. metabarcoding at family-level 19
  - Taxa unique to one approach 19
  - Family-level vs genus-level 21
- Environmental effects 22
- Compositional Turnover (beta diversity) 25


---

This supplementary file describes the inputs and model structure for the analysis of occupancy of macroinvertebrates based on composition derived from morphological identification and DNA metabarcoding. Details of the Canadian Aquatic Biomonitoring Network's wetland protocol are available from Environment and Climate Change Canada online. The metabarcoding protocols are given in the main manuscript, and detailed in *Gibson et al. 2015 PLoS One* https://doi.org/10.1371/journal.pone.0138432.

---

### Input Data

#### Sampling sites

**Table S1.1:** Coordinates of sampling sites analysed in this study (also displayed in Figure 1 in the main text).

| Site Name | Site No. | Latitude | Longitude |
|---|---|---|---|
| Otter Creek | 1 | 58.60273 | -111.5261 |
| Mamawi Bay | 3 | 58.56475 | -111.5108 |
| Mamawi Creek Pond | 4 | 58.50773 | -111.5180 |
| Child's River | 11 | 58.63840 | -111.5965 |
| Rat Lake | 14 | 58.87465 | -111.3248 |
| Egg Lake | 33 | 58.88236 | -111.3992 |
| Rocher River | 37 | 58.83234 | -111.2807 |
| Horseshoe Slough | 38 | 58.86389 | -111.5816 |

During the initial development phase of the monitoring program, between 2012 and 2014, sampling took place twice a year in June and August, and the majority of wetland sites were sampled three times each (i.e. triplicate benthic samples). However, after the initial period of funding ended, the cost of sustaining multiple field surveys and a high degree of replication was not sustainable. As a result, field surveys have since been conducted in August, and sample processing has been sustained to characterize inter-annual variability, rather than within-site replicate similarity or seasonal variability.

The samples metabarcoded in 2011 were analysed using the BR5 primer set (*Hajibabaei et al. 2011 PLoS One, Gibson et al. 2014 PNAS*) only, but thereafter both BR5 and the F230 primer set (*Gibson et al. 2015 PLoS One*) were used to process all metabarcoded samples. Please note that in the supplementary of *Gibson et al. 2014*, ArR5, the reverse primer of the BR5 primer set, was not written in reverse complement and should have been listed as follows:
5' GTRATIGCICCIGCIARIACIGG

**Table S1.2:** Number of invertebrate samples collected from the same eight sites in the Peace-Athabasca Delta in northern Alberta between 2011 and 2016, processed either using traditional morphological identification based on the standard guidelines of the Canadian Aquatic Biomonitoring Network (CABIN), or by using DNA metabarcoding.

| Year | 2011 | 2012 | 2012 | 2013 | 2013 | 2014 | 2014 | 2015 | 2016 | Total |
|---|---|---|---|---|---|---|---|---|---|---|
| Month | Aug | Jun | Aug | Jun | Aug | Jun | Aug | Aug | Aug | |
| CABIN | 8 | 24 | 24 | 24 | 8 | 8 | 8 | 12 | 10 | 126 |
| DNA | 24 | 24 | 23 | 24 | 20 | 0 | 8 | 7 | 8 | 138 |

---

#### Environmental parameters

The Peace-Athabasca is remote, making it challenging to access sites and collect data, and to leave and retrieve recording devices. The system is naturally hydrologically dynamic with a high variation in the frequency of flooding among sites, and between years. If spring snowmelt is combined with an ice-jam, water levels can rapidly increase by 2 meters or more, which is sufficient for floodwaters to connect adjacent wetlands. The potential impacts of high flow and ice are that recording devices are often damaged or lost, even at the permanent monitoring stations. Below we show where those gaps occurred in the environmental records and how we related available data to describe the variation in site condition.

---

**Temperature**

Air temperature is publicly available online from Environment Canada meteorological station at Fort Chipewyan (Gauge #71305, -111.12 W, 58.77 N). The record for water temperature at each of our study sites however was discontinuous, and there was evidence of a site level effect on their relationship to air temperature ($F$=7.079, $p$=0.0008). Annual changes in water temperature were therefore predicted separately for each site, using mean air temperature when it exceeded 0'C. We tested a range of lag periods and functions, and found water temperature could be predicted with a reasonable degree of accuracy ($r2$>0.8) using only one or two covariates: the mean temperature of the preceding 3 and 14 day period. These models were subsequently used to predict continuous water temperature profiles for each site, and estimate 3 covariates:

1. the period of time each site will have been free of ice at the time of sampling (i.e. time >0'C prior to June or August surveys).
2. the mean water temperature the month prior to sampling
3. the maximum water temperature of the year prior to sampling

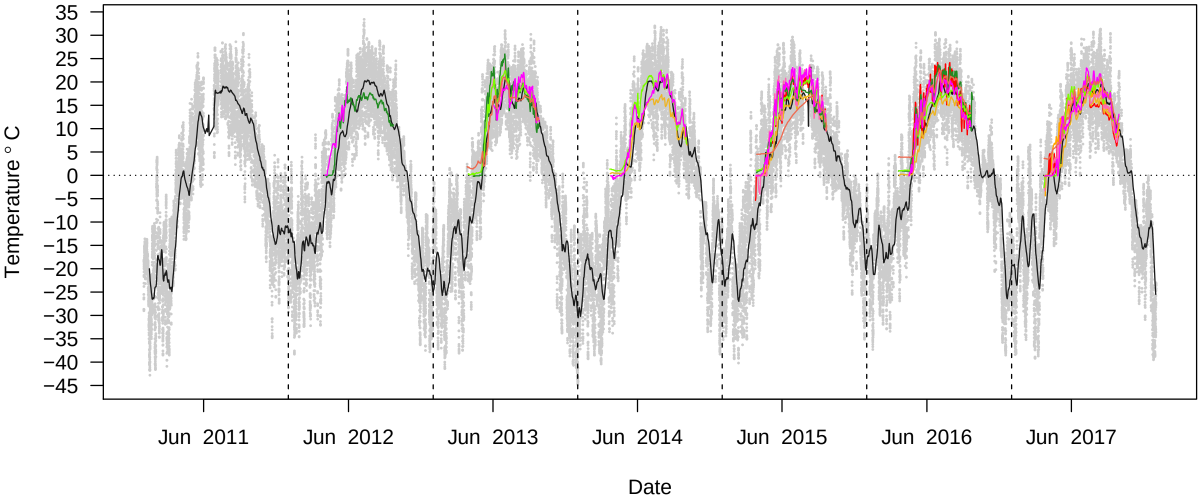


**Figure S1.1:** Range of air temperatures recorded hourly at Fort Chipewyan, adjacent to the PAD, between 2011 and 2017 (grey). This information was summarised to 3-day (not shown) and 14-day mean air temperatures (black line), and used to predict water temperature. The coloured lines indicate the periods for which we had records of water temperature at each site to fit those models.

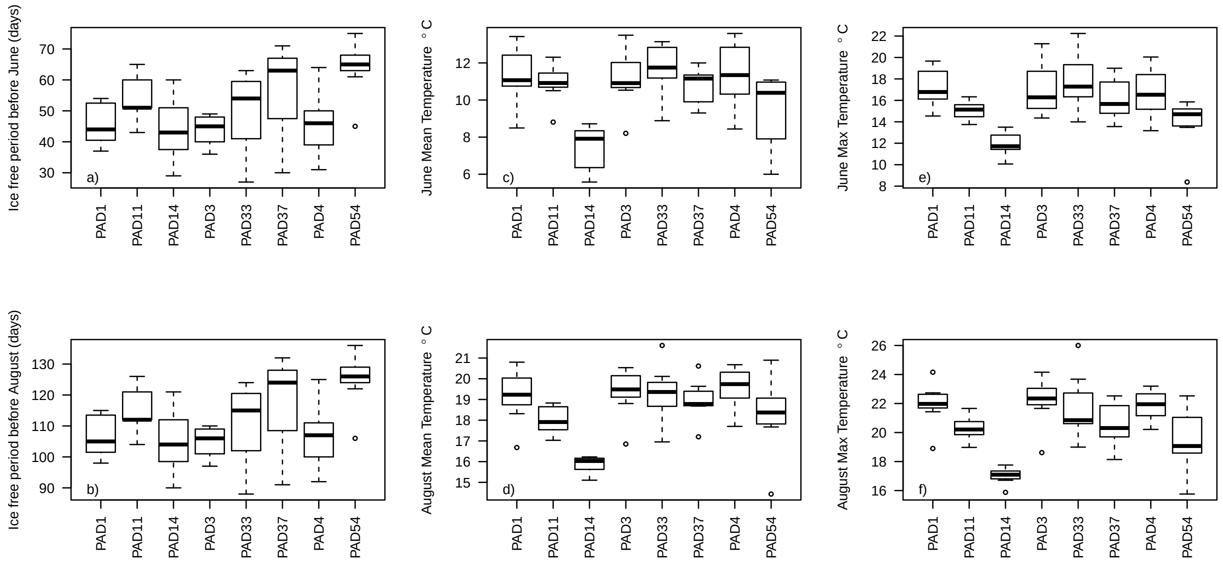


**Figure S1.2:** Site-level variation for the three covariates for water temperature regime used in our analysis divided between June and August: ice-free period (a and b), mean water temperature (c and d), and maximum water temperature (e and f).

---

**Flood frequency and duration**

Like temperature, our study made use of water level data collected by Environment and Climate Change Canada from the surveyed sites (*Peters et al. 2006 Hydrological Processes*, *Peters et al. 2016 Canadian Water Resources Journal*, see also ECCC 2018. HYDAT Water Survey Data Products available at: https://www.canada.ca/en/environment-climate-change/services/water-overview/quantity/monitoring/survey/data-products-services.html). However, we could not calibrate those discontinuous site-specific hydrological profiles against a single continuously recording station because no such reference exists. There are currently 11 gauge stations operating on either the Peace and Athabasca rivers entering the PAD, in the PAD, and on the Slave river that drains from the delta. However, during the timeline of our study, it was rare for all gauges to be recording at the same time, and importantly, records of the flood peaks were missing from some locations. Ultimately, to predict water level at specific wetlands, we first fit models to predict values for the gaps in the water level records of important gauging stations, using as inputs the values of other gauges that were still operating. A continuous estimate of water levels at each study site was subsequently based on all relevant gauging stations.

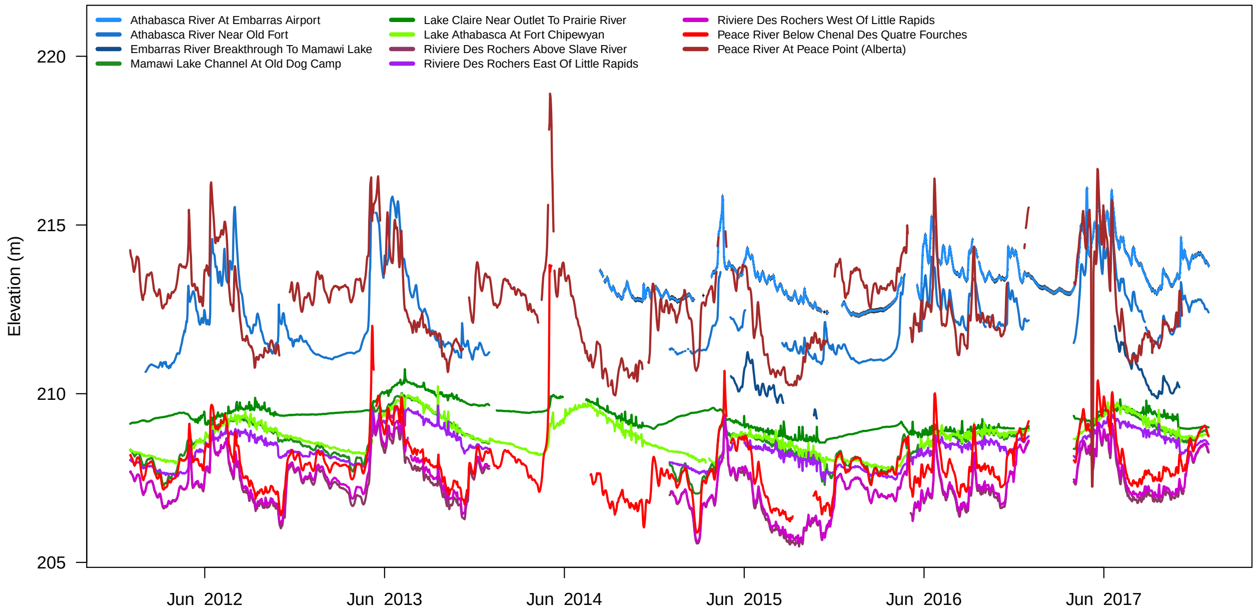


**Figure S1.3:** Water level at the 11 gauges associated with the Peace-Athabasca Delta between 2012 and 2017.

As was the case for the permanent flow gauges, there were gaps in the records available for water level in each of our study sites (Figure S1.4). It is worth noting that the delta is extensive because the landscape is very flat, and often small changes in water elevation are enough to connect main channels to adjacent wetlands. Furthermore, note that there were no measurements of water-level at these particular sites during the first season of field surveys in 2011. Although surveys in 2017 do not form part of the analysis in this study, we did take advantage of the water-level records from 2017 in order to improve our estimates of site-level hydrology.

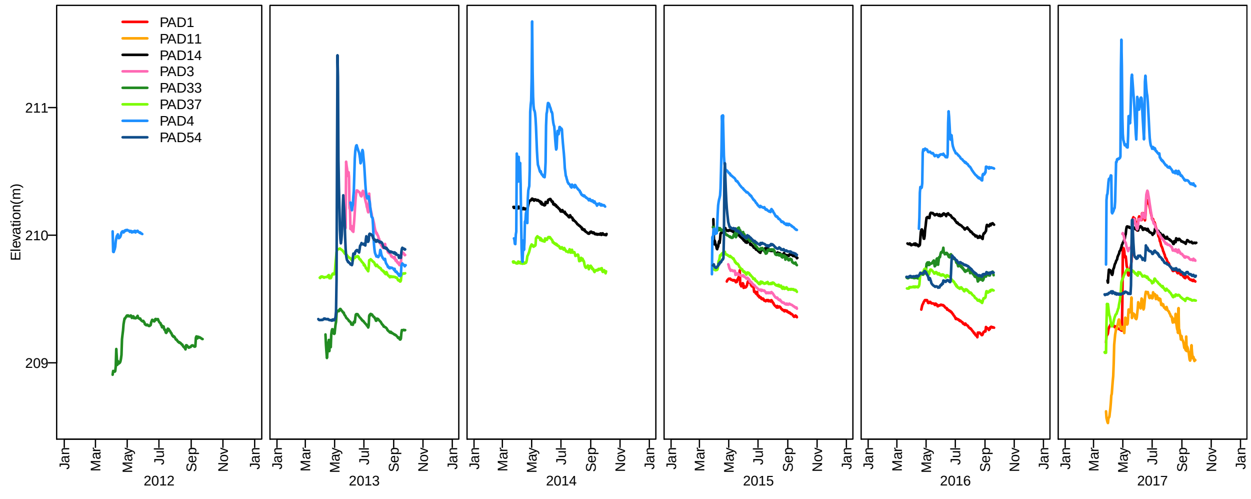


**Figure S1.4:** Record of water level from loggers deployed at the eight study sites between 2012 and 2017.

Data on the water-level at our study sites was most related to water level at gauging stations during floods, because this is when the water in the channels and adjacent wetlands was most likely connected. Thus our aim was to predict when and how often river levels exceed the wetland spill elevation, and for how long, whereas a model of the precise water level within our study site was beyond our needs. Models were calibrated to recover dynamics whose peak flows corresponded to rapid changes in water level within the wetland (Figure S1.3 and S4). We were therefore able to characterise inter-annual variation in the hydrology and connectivity of each wetland based on the frequency and duration of flooring in the spring and summer months before they were surveyed. Note the duration of time a site was flooded in a given year was positively associated with the frequency of flooding (r=0.57).

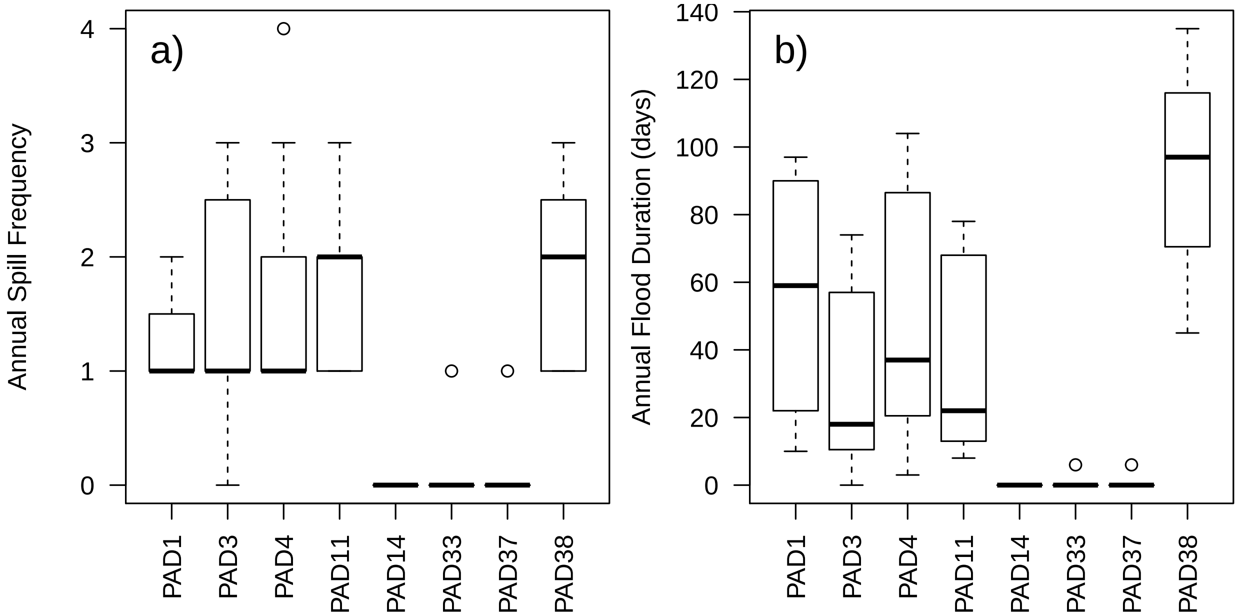


**Figure S1.5:** Variability in estimated a) annual flood frequency, and b) flood duration, per site in this study.

---

#### Assemblage Data

##### Morphological Data

Wetland invertebrate samples identified based on morphological characteristics followed standard protocols that mirror the traditional guidelines used to assess river and stream kick samples in the Canadian Aquatic Biomonitoring Network (CABIN), and hence will hereafter be referred to as the CABIN samples. The taxonomic resolution of CABIN data varies among invertebrate groups depending on the difficulty of identification, the perceived importance of a particular group and the expertise of the taxonomist. Thus although most of the 74 taxa identified were at the family-level, and subsequent analysis refers to CABIN data at the family-level, with three exceptions:
1. Information on the four sub-families of Chironomidae was retained; namely Chironominae, Orthocladiinae, Tanypodinae & Tanytarsini.
2. The Subclass Hirudinea was retained.
3. The Subclass Oligochaeta was retained.

---

##### DNA Metabarcoding Data

Raw paired-end Illumina MiSeq reads were processed using the SCVUC v 2.1 COI metabarcode pipeline available from https://github.com/Hajibabaei-Lab/SCVUC_COI_metabarcode_pipeline
Jobs were run in parallel using GNU parallel [1]. Briefly, reads were paired using SEQPREP v1.2 ensuring a minimum Phred quality of 20 and a minimum overlap of 25 bp [2]. Primers were trimmed using CUTADAPT v1.15 ensuring a minimum trimmed sequence length of 150 bp, mimimum Phred quality of 20 at the ends, and no more than 3 ambiguous bases allowed [3]. Sequences were dereplicated using VSEARCH 2.7.0 '–derep_fulllength' [4]. Unique sequences were denoised using USEARCH v10.0.240 with the unoise3 algorithm [5]. Denoising involves the removal of rare sequence clusters, potential PhiX carryover, sequences with

putative errors, and putative chimeric sequences. We defined rare sequence clusters as containing only one or two sequences. An ESV x sample table was build using VSEARCH '–usearch_global' setting the identity to 1.0 (100% similarity). Taxonomic assignments were performed using the COI Classifier v3 available from https://github.com/terrimporter/CO1Classifier [6]. This classifier uses the Ribosomal Database Project naïve Bayesian classifier with a COI reference set mined from GenBank [7]. We used the recommended cutoffs to ensure taxonomic assignments are 99% correct, assuming the query taxa are represented in the database.

Bioinformatics references:
[1] O. Tange, GNU Parallel - The Command-Line Power Tool. ;login: The USENIX Magazine February, 42-47 (2011).
[2] J. St. John, SeqPrep (2016).
[3] M. Martin, Cutadapt removes adapter sequences from high-throughput sequencing reads. EMBnet. journal 17, pp-10 (2011).
[4] T. Rognes, T. Flouri, B. Nichols, C. Quince, F. Mahé, VSEARCH: a versatile open source tool for metagenomics. PeerJ 4, e2584 (2016).
[5] R. C. Edgar, UNOISE2: improved error-correction for Illumina 16S and ITS amplicon sequencing. bioRxiv (2016) https:/doi.org/10.1101/081257 (June 28, 2018).
[6] T. M. Porter, M. Hajibabaei, Automated high throughput animal CO1 metabarcode classification. Scientific Reports 8, 4226 (2018).
[7] Q. Wang, G. M. Garrity, J. M. Tiedje, J. R. Cole, Naive Bayesian Classifier for Rapid Assignment of rRNA Sequences into the New Bacterial Taxonomy. Applied and Environmental Microbiology 73, 5261-5267 (2007).

As the sampling approach entails disturbance of emergent vegetation, it is common for wetland samples to contain a variety of invertebrates with a terrestrial, rather than freshwater origin. For the purposes of this study we focused solely on freshwater taxa, and taxa whose lifecycle was not known to include an aquatic freshwater phase were discarded. The resulting dataset included 263 invertebrate genera among 109 families, and consistent with most ecological studies, many taxa were rarely observed, and few were common. Those families and genera we could confidently identify were among the 29,099 and 15,587 unique sequences returned by the BR5 and F230 respectively. As Figure S1.6d shows, the extremely high abundance of rare sequences, and lack of sequences shared among a majority of sites, does not support an analysis of occupancy, and hence this study focused on the substantial diversity observable at the genus and family levels.

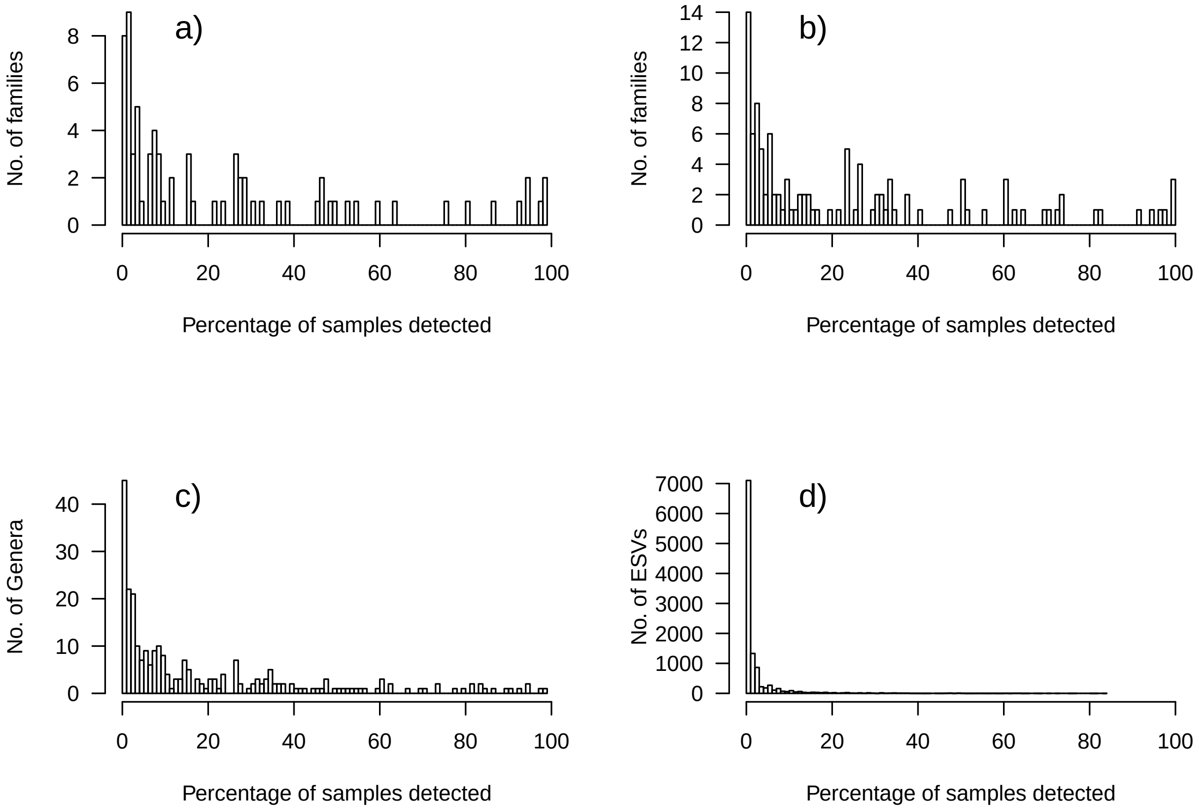


**Figure S1.6**: Prevalence of taxonomic units in the 126 (a) and 138 (b,c,d) samples processed using CABIN or DNA metabarcoding. Panel a) are morphologically identified taxa, predominantly at the family-level, b) metabarcoding at the family-level, c) metabarcoding at the genus-level, and d) Exact Sequence Variants (ESV).

---

**Exploration of community data**

Firstly, looking at the metabarcoding data at the rank of genera, we found that compositional turnover among replicate samples from the same site, at the same time, and using the same primer, was on average greater than 30%:

**Table S1.3:** Mean genus-level dissimilarity of replicate samples collected from the same site, and during the same visit, as identified by DNA metabarcoding using two sets of primers.

| | PAD1 | PAD3 | PAD4 | PAD11 | PAD14 | PAD33 | PAD37 | PAD38 | Mean |
|---|---|---|---|---|---|---|---|---|---|
| BR5 | 0.27 | 0.26 | 0.34 | 0.27 | 0.31 | 0.42 | 0.32 | 0.36 | 0.32 |
| F230 | 0.34 | 0.30 | 0.34 | 0.28 | 0.29 | 0.35 | 0.28 | 0.33 | 0.31 |

Secondly, turnover was typically greater than 50% among sites from the same survey period (again using the same primer).

**Table S1.4:** Mean genus-level dissimilarity of samples collected from different sites within the Peace-

Athabasca Delta, during the same visit (same season and year), as identified by DNA metabarcoding using two sets of primers.

| | PAD1 | PAD3 | PAD4 | PAD11 | PAD14 | PAD33 | PAD37 | PAD38 | Mean |
|---|---|---|---|---|---|---|---|---|---|
| BR5 | 0.54 | 0.50 | 0.56 | 0.54 | 0.54 | 0.50 | 0.52 | 0.54 | 0.53 |
| F230 | 0.52 | 0.48 | 0.51 | 0.51 | 0.50 | 0.47 | 0.48 | 0.50 | 0.50 |

Finally, we could expect compositional dissimilarity to gradually increase over time before reaching a plateau over longer timescales. However, relative to the rapid turnover observed among replicate samples, and among sites, dissimilarity of samples from the same location separated in time remained relatively uniform for most sites. We also did not find any evidence to suggest that taxa were any more likely to be detected at the same site in subsequent years than random, given their prevalence in our dataset.

Note that although there appeared to be a sharp increase in dissimilarity after 5 years, we only had a single pairwise comparison per site over this length of time, whereas many more combinations exist for samples taken 1-3 years apart.

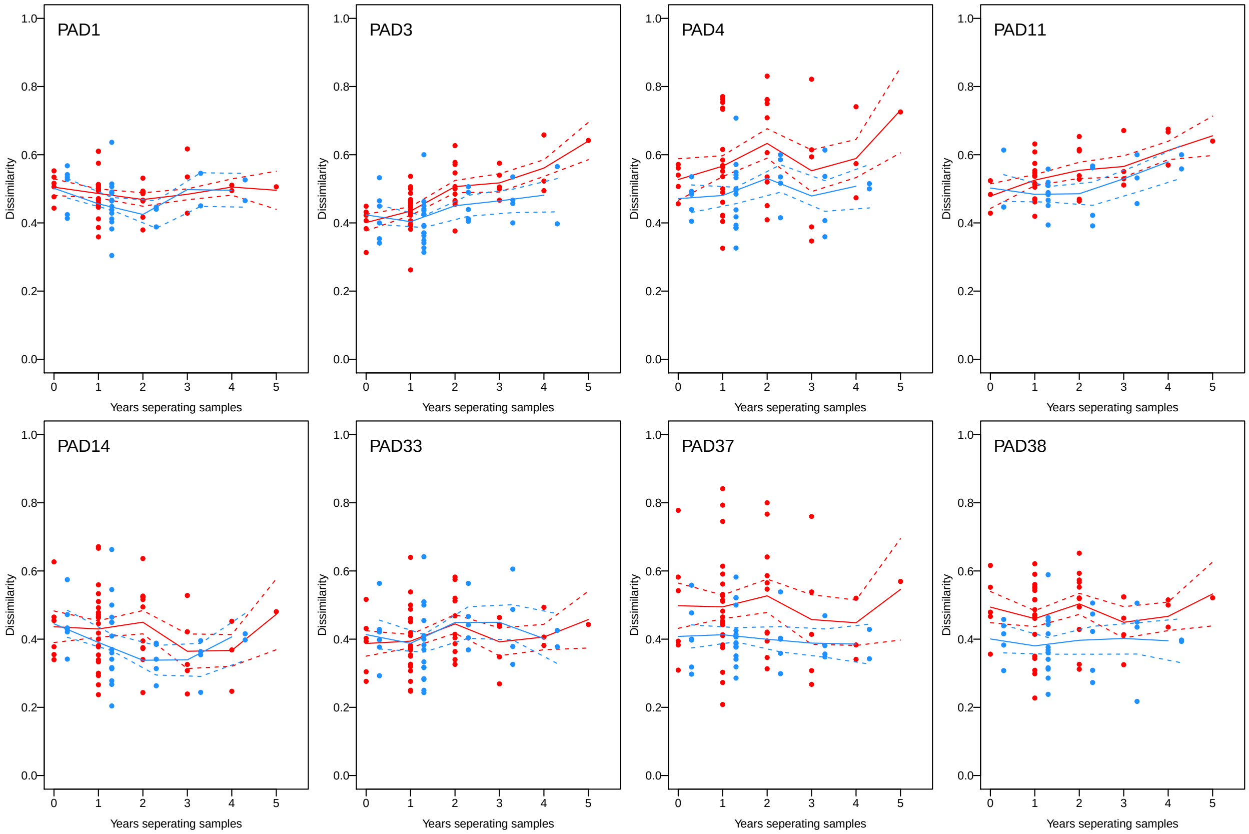


**Figure S1.7:** Relationship between invertebrate pairwise dissimilarity (Sorenson index) from the same site location given their seperation in years. See Table S1.1 for locations based on site numbers. Dissimilarity was calculated for both BR5 (red) and F230 composition (blue), and lines indicate the mean and 95% confidence intervals of a polynomial regression.

### Hierarchical Model Framework

#### Occupancy Modelling

This study followed the framework and model structure outlined by Kery and Royle (*2015, Applied Hierarchical Modeling in Ecology: Analysis of distribution, abundance and species richness in R and BUGS: Volume 1*) and we urge interested readers to consult this text for further information.

As described in the main text, four models were constructed to represent the various data types avaiable:
1. *CABIN Fcount*: CABIN counts of families
1. *CABIN Fpa*: CABIN presence-absence of families
1. *DNA Fpa*: DNA presence-absence of families
1. *DNA Gpa*: DNA presence-absence of genera

*Detection Matrix*
The community composition matrices were formatted as arrays for sites, replicates and taxa. The dimension *sites* refers to the 62 unique combinations of 8 sites, 1 or 2 seasons, and 6 years. The *replicates* dimension includes 3-6 columns based on a maximum number of attempts to observe the composition of the community during site visits. For CABIN sampling, triplicate samples were equivalent to 3 observations, but for metabarcoding, the processing of samples twice with separate pairs of primers represents independent observations of the community, and hence up to 6 replicate observations. Detection of all *taxa* is then reflected in each layer of the array, namely 74 and 109 at family-level for CABIN and metabarcoding respectively, and 263 at the genus-level for metabarcoding. Note that where there was no survey e.g. because we did not survey all sites in triplicate every year, the entries in the detection matrix were set as NA's.

*Data Augmentation*
To extend the model infernce to encompass taxa that were never detected requires a process known as data-augmentation (see *Dorazio and Royle, 2006 Ecology*, and more recently *Guillera-Arroita 2019 Ecology and Evolution*). The key addition to the basic occupancy model structure is an overarching process that determines how many taxa from a larger superpopulation $M$ are expected to be included in the system. The fraction of $M$ that are included is described by $\Omega$. Ideally the parameter $M$ would be informed by some prior knowledge of how many species were present in the broader region but given we do not know this, we chose a vague prior, large enough that the posterior distribution of $\Omega$ was not close to 1, but also not so high that the model took far longer to converge.

Thus, the detection matrix was expanded to allow for data augmentation. See page 682 of *Kery and Royle 2015* for more details:

```
# Total number of Genera identified by metabarcoding
nspec
# 263
# Vague prior on Ntotal: M = approximately 50% more than nspec
nz   = (400-nspec)
# Empty array with only zeroes
Yaug = array(0, dim=c(nsite, nrep, nspec+nz))
# Copy into it the observed data
Yaug[,,1:nspec] = Y
# Now create same NA pattern in the augmented matrix, as was the case for the observed taxa.
missings = is.na(Yaug[,,1])                         # e.g. no replicates in 2016
for(k in (nspec+1):(nspec+nz)){ Yaug[,,k][missings] = NA }
```

---

##### JAGS Models

The code below provides the structure for the occupancy-detection model, with numbers taken from the DNA metabarcoded genus-level model, but the format was the same for the family-level models too. For illustration, the code includes indicator variables, which are used to estimate the importance of a given covariate in model fitting, and although the starting model included 7 possible covariates, the code below has been reduced to three.

A reminder of the inputs:

```
# Bundle and summarize data
win.data = list(Y      = Yaug,              # Response data
                nsite  = dim(Y)[1],         # No. of sites
                nrep   = dim(Y)[2],         # Max no. of replicates
                nspec  = dim(Y)[3],         # No. of observed species
                nz     = nz,                # No. of hypothetical species
                M      = nspec + nz,        # Metacommunity richness
                X      = X,                 # Environmental covariates
                ncov   = ncol(X)            # No. of covariates
)

# Specify model in BUGS language
sink("PAD_Gen_Occ_model.txt")
cat("
    model {

    # Data Augmentation priors - Fraction of superpopulation present in this community
    omega ~ dunif(0,1)

    # Priors for species-specific effects in occupancy and detection
    for(k in 1:M){
      # Occupancy intercept
      lpsi[k]       ~ dnorm(mu.lpsi, tau.lpsi)
      # Occupancy response slope
      for(i in 1:ncov){
        betalpsi[k,i]   ~ dnorm(mu.betalpsi[i], tau.betalpsi[i])
      }
      # Detectability (seperate estimate per primer pair)
      for(j in 1:2){
        lp[k,j]         ~ dnorm(mu.lp[j], tau.lp[j])
      }
    }

    # Hyperpriors

    # For the model of occupancy
    mu.lpsi         ~ dnorm(0,0.1)               # Community mean of occupancy
    tau.lpsi        = pow(sd.lpsi, -2)
    sd.lpsi         ~ dunif(0,8)                 # Standard deviation of occupancy

    for(i in 1:ncov){
      mu.betalpsi[i]    ~ dnorm(0,0.1)           # Occupancy response to each covariate
      tau.betalpsi[i]   = pow(sd.betalpsi[i], -2)
      sd.betalpsi[i]    ~ dunif(0, 4)            # Standard deviation of coefficent
    }

    # Indicator variables (used when selecting which covariates to retain).
    for(i in 1:ncov){
      V[i] ~ dbern(0.5)
```

```
    }

    # Community mean(s) of detection (logit) divided by the type of Primer
    for(j in 1:2){
      mu.lp[j]           ~ dnorm(0,0.1)
      tau.lp[j]          = pow(sd.lp[j], -2)
      sd.lp[j]           ~ dunif(0, 4)
    }

    # Data Augmentation process: Ntotal taxa sampled out of M available
    for(k in 1:M){
      w[k] ~ dbern(omega)
    }

    # Ecological model for true occurrence (process model)
    for(k in 1:M){
      for(i in 1:nsite) {
        logit(psi[i,k]) = lpsi[k] +
                          (betalpsi[k,1] * X[i,1] * V[1]) +
                          (betalpsi[k,2] * X[i,2] * V[2]) +
                          (betalpsi[k,3] * X[i,3] * V[3])
        mu.psi[i,k]     = w[k] * psi[i,k]
        z[i,k]          ~ dbern(mu.psi[i,k])
      }
    }

    # Observation model for replicated detection/nondetection observations
    for(k in 1:M){
      for(i in 1:nsite){
        for(j in 1:nrep){
          logit(p[i,j,k]) = lp[k,PRI[j]]
          mu.p[i,j,k]     = z[i,k] * p[i,j,k] * step(anyseq[i,j]-1)
          Y[i,j,k]        ~ dbern(mu.p[i,j,k])
        }
      }
    }

    # Derived quantities
    for (i in 1:nsite){ Nsite[i] = sum(z[i,]) }     # Number of species at each site
    n0      = sum(w[(nspec+1):(nspec+nz)])          # Number of unseen species
    Ntotal  = sum(w[])                              # Total metacommunity size

    # Vectors to save (capital S in name is for 'save')
    # To save space, discard samples for all except 1 of the potential species.
    lpsiS[1:(nspec+1)]            = lpsi[1:(nspec+1)]
    betalpsiS[1:(nspec+1),1:ncov] = betalpsi[1:(nspec+1),1:ncov]
    lpS[1:(nspec+1),1:2]          = lp[1:(nspec+1),1:2]

    }
    ",fill = TRUE)
sink()

# Initial values
```

```r
wst = rep(1, nspec+nz)                      # Start by including every taxon
zst = array(1, dim = c(nsite, nspec+nz)) # Set every taxon to start as occurring
inits = function() list(z         = zst,
                        w         = wst,
                        lpsi      = rnorm(n = nspec+nz, sd=0.1),
                        betalpsi  = matrix(rnorm((nspec+nz)*ncol(X), sd=0.1),
                                           nrow=nspec+nz, ncol=ncol(X) ),
                        lp        = matrix(rnorm(n = (nspec+nz)*2, sd=0.1),(nspec+nz),2),
                        V         = rep(1,ncol(X)) )

# Set 1
params1 = c("omega", "mu.lpsi", "sd.lpsi", "mu.betalpsi", "sd.betalpsi",
            "mu.lp", "sd.lp", "Ntotal", "Nsite", "lpsiS", "betalpsiS","lpS","V")

# MCMC settings
ni = 1000000
nt = 50
nb = 250000
nc = 4

# Run JAGS with indicator variables
mod = jags(data = win.data,
           inits,
           parameters.to.save = params1,
           model.file = "PAD_Gen_Occ_model.txt",
           n.chains   = nc,
           n.thin     = nt,
           n.iter     = ni,
           n.burnin   = nb,
           parallel   = FALSE,
           codaOnly   = TRUE,
           verbose    = TRUE)
```

---

##### Convergence, model fit and variable selection

The most complex models included hundreds of taxa, data-augmentation and multiple-covariates, and hence convergence of all parameters was slow, evaluated based on their *Rhat* values (*Gelman and Rubin 1992 Statistical Science*). Updating initial values with previous runs did reduce run time, but ultimately some coefficients, particularly for some rare species, never fully converged. Recently, Warton et al. (*2017, Methods in Ecology and Evolution*) described how commonly used measures of model fit, particularly the Pearson chi-square goodness-of-fit statistic and Bayesian P-value (*MacKenzie and Bailey 2004, Royle et al. 2007*) were not suitable for assessing incidence-based occupancy models. We therefore followed their approach and looked at the diagnostic plots based on Dunn-Smyth residuals to assess whether different variable combinations could improve model fit.

To select which variables contribute to the occupancy model most effectively we used the latent indicator variable approach (*Kuo and Laiick 1998*). Latent variable indicators were particularly slow to converge, but to allow for potential interactions, we followed a process of backwards selection, eliminating variables if there was clear evidence they were rarely included in the fitted models once they had converged (i.e. posterior distribution of v $< 0.5$).

---

##### N-mixture Model

Occupancy models are incidence-based, modelling the detection/non-detection of each taxon. This is in part because the sequence data generated by metabarcoding is typically not regarded as quantitative, and we therefore do not have estimates of relative abundance (see *Bush et al. 2019 in press* and references therein). Conversely the processing of CABIN samples includes counts of each taxon observed in a subset of the whole sample. Given the impact of environmental changes on the aquatic invertebrate community are reflected in changes to taxon abundances, a separate "abundance-based" model, known as N-mixture modelling, was applied to the CABIN samples (*Yamaura et al. 2011 J. Applied Ecology*) to incorporate as much of the information present in CABIN samples as possible.

The underlying process of an N-mixture model assumes abundances follow a Poisson distribution, but the model structure is otherwise similar to that of the incidence-based occupancy model. The proportion of the sample processed was retained as a covariate (*alpha*) on detectability in the N-mixture model (i.e. taxon detectability and abundance was dependent on the proportion of a sample that is sorted). Note that the N-mixture model estimates the probability a given number of **individuals** were present in a sample, and the probability of detection was also estimated at the individual-level. As a result we cannot easily compare between the occupancy and N-mixture models without making some assumptions about how many individuals we can count at a site. Therefore, the intention of the N-mixture model was to compare the broad distribution of occurrence vs. detectability (Fig. S10), and the relative influence of environmental factors on those distributions.

```
# Standardize the proportion of a sample processed
# Typically 5% but ranges from 3% to 100% of the sample collected.
sort.count = standardize(perc_sorted)
# Split into site-x-replicate matrix
scount = matrix(0,ncol=3,nrow=dim(YC)[1])
for(i in 1:length(scount)){
  scount[ match( sample.list[i], dimnames(YC)[[1]] ),
          match( rep.list[i],    dimnames(YC)[[2]]) ] = sort.count[i]
}

#-x-x-x-x-x-x-x-x-x-x-x-x-x-x-x-x-x-x-x-x-x-#

# N-mixture model in BUGS language
cat("
    model {

    # Data Augmentation priors - Fraction of superpopulation present in this community
    omega ~ dunif(0,1)

    # Priors for species-specific effects in occupancy and detection
    for(k in 1:M){
      # Occupancy intercept
      lpsi[k]  ~ dnorm(mu.lpsi, tau.lpsi)
      # Occupancy response slope
      for(i in 1:ncov){ betalpsi[k,i] ~ dnorm(mu.betalpsi[i], tau.betalpsi[i]) }
      # Detection intercepts
      lp[k]         ~ dnorm(mu.lp, tau.lp)
      # Slope of detection
      alpha[k]      ~ dnorm(mu.alpha, tau.alpha)
    }

    # Hyperpriors
```

```
    # For the model of occupancy
    mu.lpsi    ~ dnorm(0,0.01)                      # Community mean of occupancy
    tau.lpsi   = pow(sd.lpsi, -2)
    sd.lpsi    ~ dunif(0,8)                         # Standard deviation of occupancy

    for(i in 1:ncov){
      mu.betalpsi[i]    ~ dnorm(0,0.1)              # Occupancy response to each covariate
      tau.betalpsi[i]   = pow(sd.betalpsi[i], -2)
      sd.betalpsi[i]    ~ dunif(0, 4)               # Standard deviation of coefficent
    }

    # Indicator variables (used when selecting which covariates to retain).
    for(i in 1:ncov){
      V[i] ~ dbern(0.5)
    }

    # Community mean(s) of detection (logit) divided by the type of Primer
    mu.lp           ~ dnorm(0,0.1)                  # Community mean of detection
    tau.lp          = pow(sd.lp, -2)
    sd.lp           ~ dunif(0, 4)

    mu.alpha        ~ dunif(-0.5,0.5)               # Mean coefficient upon detection
    tau.alpha       = pow(sd.alpha, -2)
    sd.alpha        ~ dunif(0, 4)

    # Data Augmentation process: Ntotal species sampled out of M available
    for(k in 1:M){
      w[k]          ~ dbern(omega)
    }

    # Ecological model for true occurrence (process model)
    for(k in 1:M){
      for(i in 1:nsite) {
        N[i,k]      ~ dpois(lambda[i,k] )* w[k])
        log(lambda[i,k]) = lpsi[k] +
                           (betalpsi[k,1] * X[i,1] * V[1]) +
                           (betalpsi[k,2] * X[i,2] * V[2]) +
                           (betalpsi[k,3] * X[i,3] * V[3])
      }
    }

    # Observation model for replicated detection/nondetection observations
    for(k in 1:M){
      for(i in 1:nsite){
        for(j in 1:nrep){
          logit(p[i,j,k]) = lp[k] + (alpha[k] * scount[i,j]) # step(anycount[i,j]-1)
          YC[i,j,k] ~ dbin(p[i,j,k], N[i,k])
        }
      }
    }


}"
```

### Results

#### Metacommunity size (gamma diversity)

Given the probability of detecting the taxa we observed in each dataset, data augmentation suggests how many more taxa are likely to be present in the metacommunity, but have not been observed so far. The models suggest that another ~20 families, or 100 genera, may potentially occur in the PAD system that have not been detected. Note that while data augmentation was present in each model, and hence will have to some extent influenced the overarching distribution of parameters at the community-level, the results for the augmented taxa were not stored after the model run. As a result, all the results we present below are solely for the detected taxa.

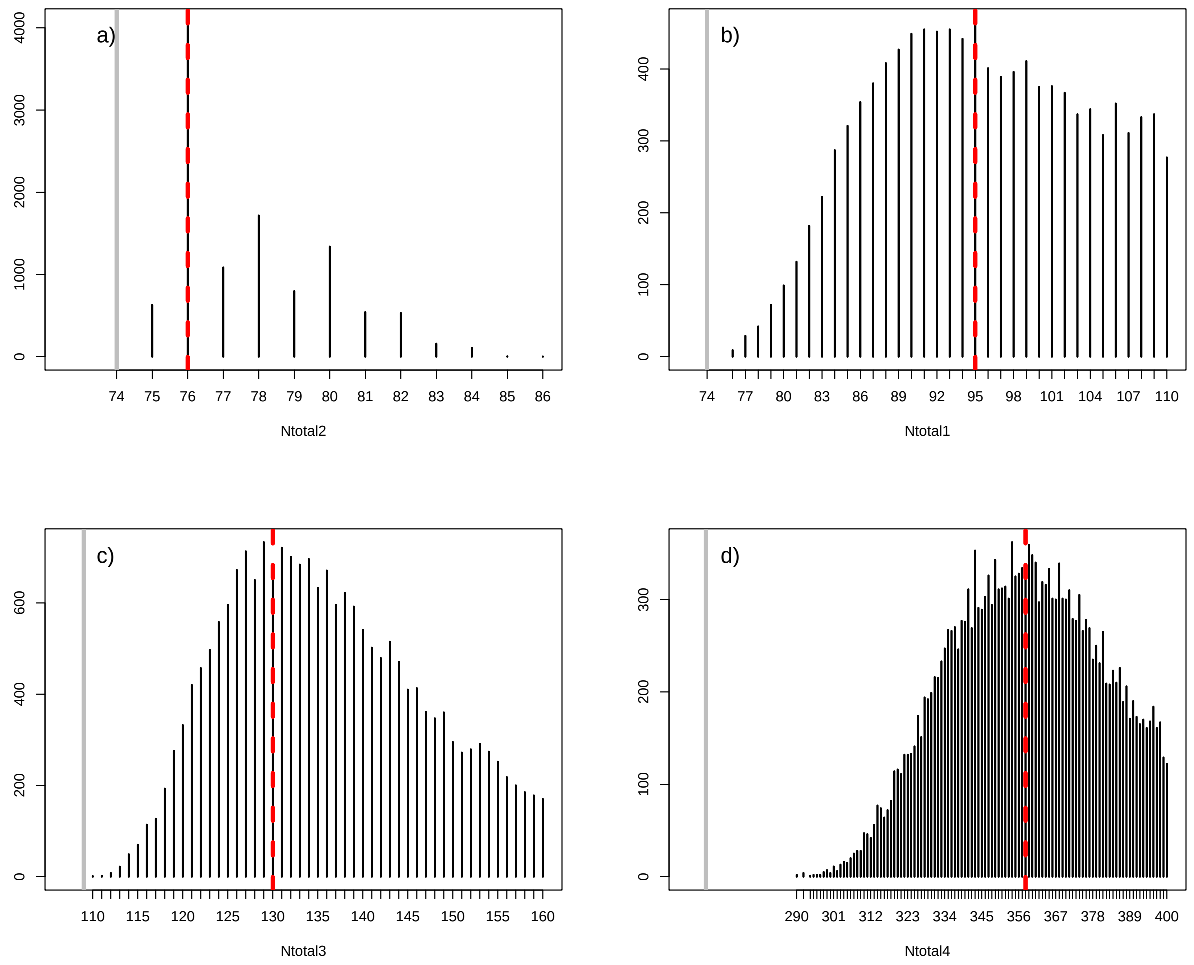


**Figure S1.8:** Estimates of the total number of taxa in the PAD metacommunity based on a) CABIN count data, b) CABIN presence/absence data, and metabarcoding data at c) family and d) genus level. Grey bars indicate the observed number of taxa, and the red dashed lines marks the mean model estimate after accounting for imperfect detection.

#### Site richness (alpha diversity)

As with the metacommunity, given the probability of detecting taxa is <1, the true richness of each sampling site is likely to be greater than what was actually observed given the sampling effort expended. The degree to which richness is underestimated varies among each model, but in every case, the true number will be greater than was observed, and Figure S1.9 simply illustrates this for the 8 sites sampled in 2016.

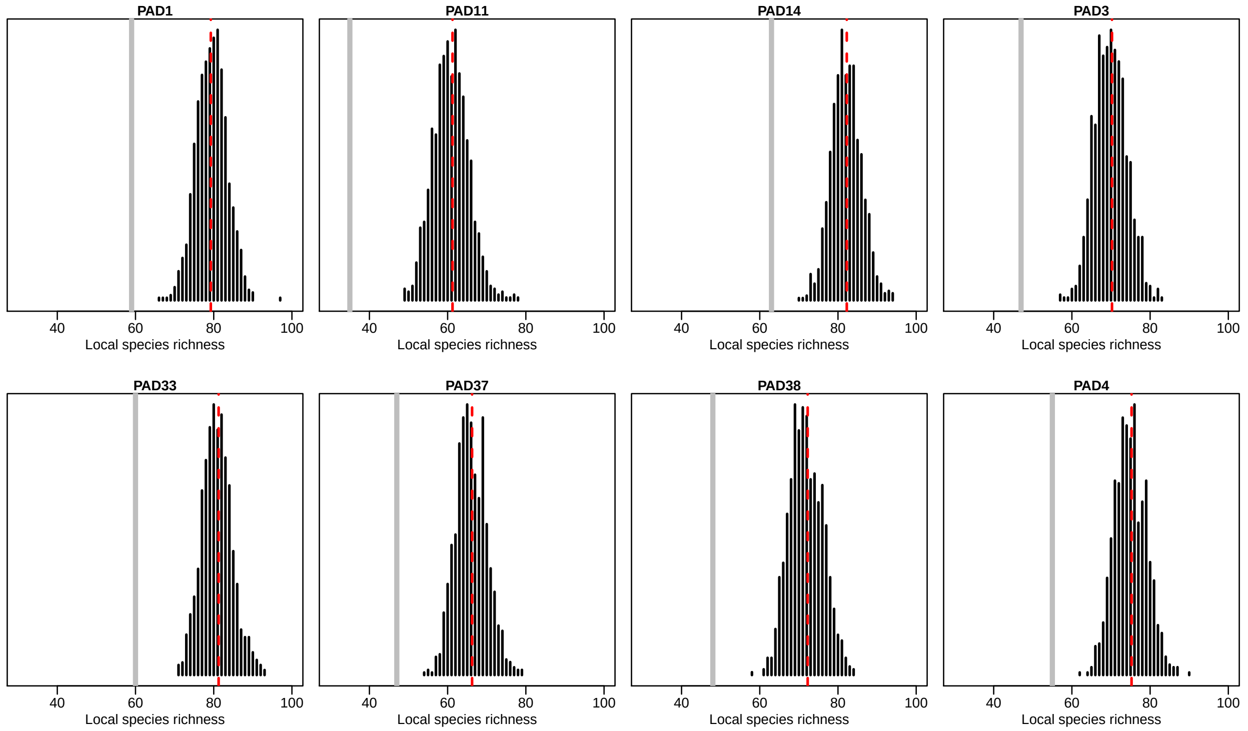


**Figure S1.9:** Estimated number of genera in each of the eight field sites during August 2016 predicted by the occupancy model trained on metabarcoding data. The grey line indicates the number of genera directly detected, and the red line marks the mean model estimate after accounting for imperfect detection.

#### Occupancy and detectability

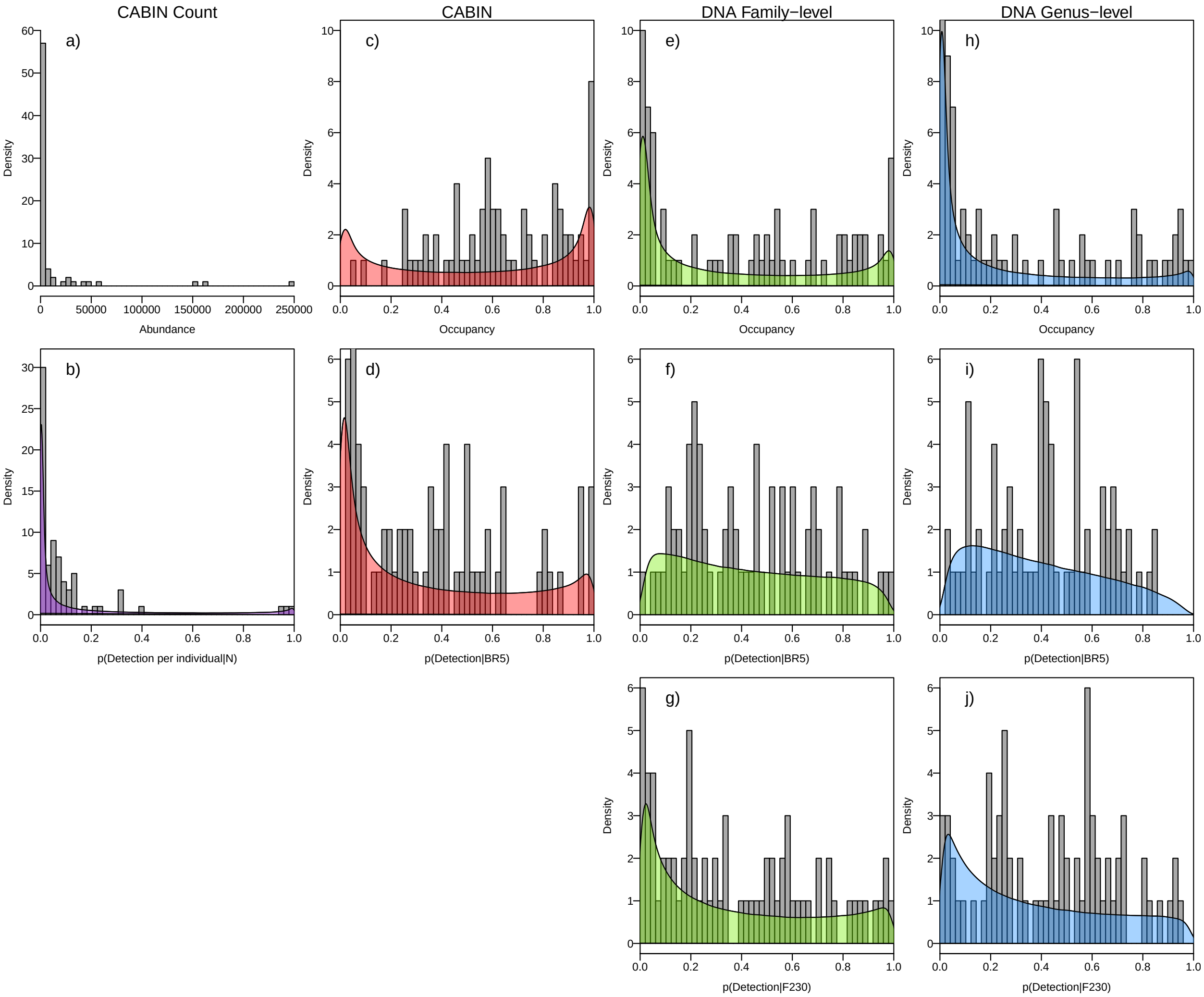


**Figure S1.10:** This figure expands upon Figure 2 in the main text to include the modelled distribution of occupancy and detectability for the CABIN N-mixture model of abundance and metabarcoding model at the genus-level. Columns show the predicted abundance and *individual* detectability based on CABIN count data (a,b), and then taxon occupancy and detectability based on CABIN presence-absence data (c,d), metabarcoding at family-level (e-g), and metabarcoding at the genus-level (h-j). Detectability of taxa with metabarcoding is subdivided into the primer pairs used by this study. The shaded polygons describe the probability density of the community-level hyperparameters, and the grey bars indicate the underlying frequency of the parameters estimated for each taxon individually.

#### CABIN vs. metabarcoding at family-level

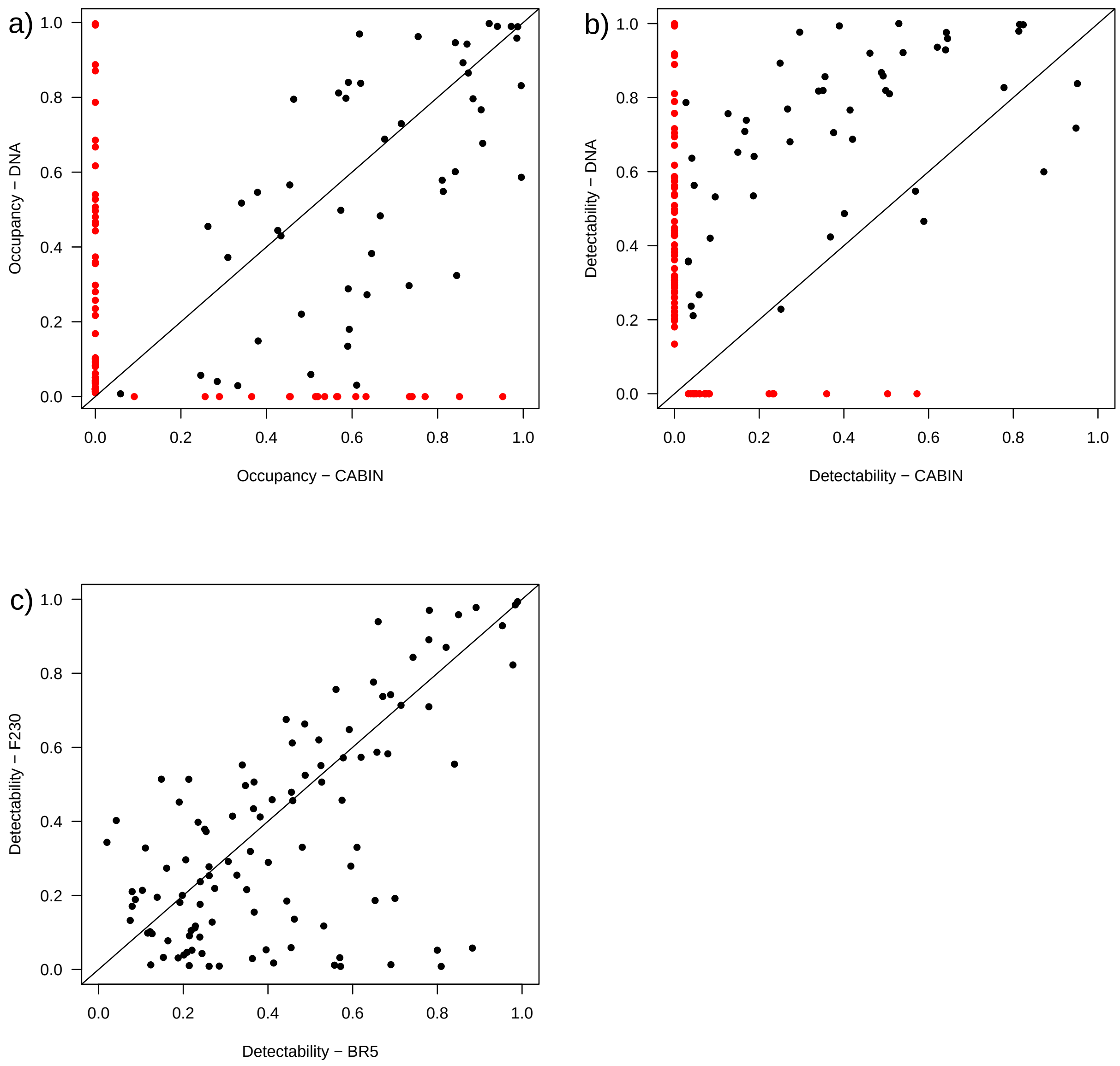


**Figure S1.11:** Comparison of a) occupancy and b) detectability estimates in models trained by CABIN data and DNA metabarcoded data at the family-level (n= 50). Red points indicate the taxa not observed by the complementary method i.e. 18 and 59 families were unique to CABIN and metabarcoding respectively. See below for further information on the identity of unique taxa. Panel c) breaks down the detectability of families detected using within DNA metabarcoding (y-axis in panel b) for each of the two primers used.

---

#### Taxa unique to one approach

Before proceeding with any comparisons, we highlight the discrepancies in the taxonomic lists between the CABIN (morphological) and metabarcoding datasets. Note that without a control sample to validate either approach (i.e. a mock community of known composition), there are several reasons why taxa may be missed by one approach or the other, and the reasons may differ among taxa. Nonetheless, we show where such discrepancies exist in our data, and explore some plausible reasons for those. We refer users to papers such as Hajibabaei et al. (*2011 PLoS ONE*), Elbrecht et al. (*2017 Ecology & Evolution*), and Bush et al. (*2019, in press*) for more discussion of comparisons between morphological and metabarcoding approaches.

Table S1.5 summarises the taxa identified by morphology in the CABIN samples, but absent from the metabarcoding dataset, and Table S1.6 provides the equivalent for taxa only identified by metabarcoding. To try to understand why some taxa might have been missed by metabarcoding, we have also included columns in Table S1.5 describing how often the taxa were detected, and the number of reference sequences available in the DNA barcode library. The first point is that the majority of taxa in Table 5 are aquatic mites, which are challenging to identify reliably, and as a result are poorly represented in DNA barcode libraries. Other taxa like Dixidae are also poorly represented in the reference library and hence the bioinformatic pipeline was less likely to relate observed sequences to these taxonomic groups. There are reasonably large numbers of reference sequences for Trichoceridae, Perlodidae, Rhyacophilidae, but given these taxa were observed in less than 10% of samples, it is possible they were simply absent from the samples processed by metabarcoding, or at such low biomass that they were less likely to be detected (see references above). Therefore, the area we would suggest is most interesting is the absence of Chaoboridae and Sminthuridae, both of which are relatively prevalent and have reference sequences (see also *Curry et al. 2018 Freshwater Science*). These specimens in the PAD may have been incorrectly identified, or reference sequences are from species within the same taxonomic group, but relatively unrelated to the species in the PAD, making assignment less certain.

**Table S1.5:** Taxonomic classification of taxa that were observed in the CABIN morphological dataset, but never observed using DNA metabarcoding. The number of reference sequences describe the coverage of each family within the library used by this study (see details in section above on metabarcoding data).

| Phylum | Class | Order | Family | No._of_samples_observed | No._of_reference_seqs |
|---|---|---|---|---|---|
| Arthropoda | Arachnidae | Trombidiformes | Mideopsidae | 1 | NA |
| Arthropoda | Insecta | Trichoptera | Uenoidae | 1 | 67 |
| Arthropoda | Insecta | Diptera | Trichoceridae | 1 | 153 |
| Arthropoda | Arachnidae | Trombidiformes | Eylaidae | 2 | NA |
| Arthropoda | Arachnidae | Trombidiformes | Sperchontidae | 2 | NA |
| Arthropoda | Arachnidae | Trombidiformes | Torrenticolidae | 2 | NA |
| Mollusca | Gastropoda | Basommatophora | Ancylidae | 2 | 48 |
| Arthropoda | Insecta | Plecoptera | Perlodidae | 3 | 271 |
| Arthropoda | Arachnidae | Trombidiformes | Hydryphantidae | 4 | 3 |
| Arthropoda | Insecta | Hemiptera | Mesoveliidae | 5 | 52 |
| Arthropoda | Insecta | Trichoptera | Rhyacophilidae | 5 | 837 |
| Arthropoda | Insecta | Diptera | Dixidae | 7 | 3 |
| Arthropoda | Arachnidae | Trombidiformes | Hygrobatidae | 10 | 144 |
| Arthropoda | Arachnidae | Trombidiformes | Oxidae | 11 | NA |
| Arthropoda | Arachnidae | Trombidiformes | Hydrodromidae | 14 | NA |
| Arthropoda | Insecta | Diptera | Chaoboridae | 45 | 95 |
| Arthropoda | Collembola | Collembola | Sminthuridae | 61 | 201 |

**Table S1.6:** Families of aquatic invertebrates included in this study that were detected using DNA metabarcoding, but were otherwise not observed using standard morphological approaches, and the percentage of samples in which they were detected using each primer set.

| Taxon | BR5_percent | F230_percent | ______ | Taxon | BR5_percent | F230_percent |
|---|---|---|---|---|---|---|
| Apatelodidae | 0.0 | 0.7 | –\|– | Isonychiidae | 0.7 | 0.0 |
| Asplanchnidae | 13.9 | 0.0 | –\|– | Lecanidae | 6.6 | 5.1 |
| Belostomatidae | 1.5 | 0.7 | –\|– | Lepadellidae | 8.8 | 0.0 |
| Bosminidae | 4.4 | 0.0 | –\|– | Leptodoridae | 0.7 | 0.0 |

| Taxon | BR5_percent | F230_percent | ______ | Taxon | BR5_percent | F230_percent |
|---|---|---|---|---|---|---|
| Brachionidae | 3.6 | 2.9 | –\|– | Limoniidae | 2.2 | 2.9 |
| Brachyceridae | 2.2 | 6.6 | –\|– | Lubomirskiidae | 1.5 | 14.6 |
| Caeciliusidae | 2.2 | 2.2 | –\|– | Lumbricidae | 0.0 | 0.7 |
| Candonidae | 16.8 | 25.5 | –\|– | Lumbriculidae | 45.3 | 42.3 |
| Chaetonotidae | 0.7 | 2.9 | –\|– | Metropodidae | 0.7 | 0.7 |
| Chloropidae | 2.9 | 1.5 | –\|– | Muscidae | 0.7 | 0.0 |
| Chrysopidae | 0.7 | 0.0 | –\|– | Noctuidae | 0.7 | 1.5 |
| Chydoridae | 59.1 | 13.1 | –\|– | Perlidae | 1.5 | 0.0 |
| Cicadellidae | 3.6 | 0.0 | –\|– | Plumatellidae | 18.2 | 10.9 |
| Corycaeidae | 0.7 | 0.0 | –\|– | Poduridae | 2.2 | 5.1 |
| Cyclopidae | 26.3 | 0.0 | –\|– | Polyphemidae | 34.3 | 0.0 |
| Cyprididae | 78.1 | 81.0 | –\|– | Scathophagidae | 2.9 | 2.9 |
| Daphniidae | 99.3 | 69.3 | –\|– | Scirtidae | 2.9 | 4.4 |
| Diaptomidae | 54.0 | 51.8 | –\|– | Sialidae | 0.7 | 0.0 |
| Dolichopodidae | 3.6 | 1.5 | –\|– | Sididae | 26.3 | 5.1 |
| Elmidae | 1.5 | 0.0 | –\|– | Simuliidae | 7.3 | 0.7 |
| Enchytraeidae | 0.7 | 0.0 | –\|– | Sisyridae | 2.2 | 5.8 |
| Erpobdellidae | 19.0 | 13.9 | –\|– | Spongillidae | 11.7 | 0.0 |
| Eulophidae | 0.7 | 0.7 | –\|– | Stenostomidae | 13.1 | 0.0 |
| Eurycercidae | 74.5 | 4.4 | –\|– | Synchaetidae | 0.7 | 0.7 |
| Glossiphoniidae | 13.9 | 29.2 | –\|– | Tabanidae | 1.5 | 2.9 |
| Helophoridae | 0.7 | 0.7 | –\|– | Trichocercidae | 2.2 | 0.0 |
| Hydraenidae | 0.7 | 1.5 | –\|– | Tubificidae | 100.0 | 82.5 |
| Hydropsychidae | 3.6 | 0.0 | –\|– | Ulidiidae | 0.7 | 0.0 |
| Hypsibiidae | 13.9 | 9.5 | –\|– | NA | NA | NA |

---

**Family-level vs genus-level**

Figure S1.12 compares the estimates of occupancy and detectability derived from the *DNA Fpa* and *DNA Gpa* models. The majority of genera are less prevalent than their parent ranks. Naturally single-genus families are as common and hence lie on the 1:1 line, but for groups with multiple genera, occupancy could be much lower (e.g. see Chironnomidae in red), and those differences in occupancy may reveal difference sin environmental preferences and potentially competitive interactions. If the most common genus in a family (e.g. Culicidae and Glossiphonidae) was not as common as the parent-family, the distributions of genera were at least partially non-overlapping, and hence there may be evidence to suggest these differences are the reuslt of niche differences or competition. Likewise, rare genera may coexist with more common members of the family, but their narrow range may again be the result of niche differences.
With respect to detectability, we would expect specific genera to be more difficult to observe than families simply because they are less abundant than all members of the family combined. Nonetheless, there are instances when detectability of a specific genus is actually much higher than at the family-level because estimates were averaged for other genera. Note that although genera cannot be more common than their parent-families, estimates of occupancy are conditional on the likelihood of detection, and in some cases the *DNA Gpa* predicted occupacy marginally higher than that of the *DNA Fpa.* This is likely to occur if a genus was observed in many locations, and was actually quite hard to detect.

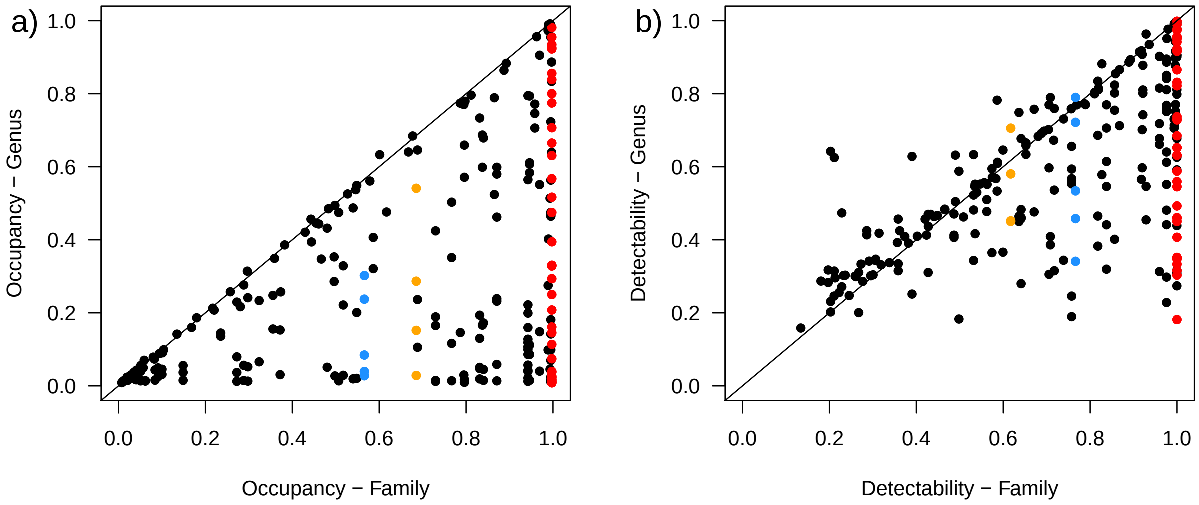


**Figure S1.12:** Comparison of a) occupancy and b) detectability between models at the family-level and genus-level. Chironomidae (red), Culicidae (blue) and Glossiphonidae (orange) are highlighted to illustrate the variation among genera.

---

#### Environmental effects

Based on the latent indicator variables for the occupancy and N-mixture models based on CABIN data, environmental covariates were included in less than 0.3% of saved iterations.

Occupancy models based on metabarcoding data identified relationships between the occupancy of taxa and environmental factors, but these were generally weak, with the response of most taxa not significantly different from 0. The indicator variables consistently identified the frequency of flooding, the length of time the site had been free of ice, and the maximum temperature that year for inclusion. Figure S1.13 illustrates the specific responses of each genus to each covariate, and the overarching weak effect at the community level. Figure S1.14 further identifies for which taxa these effects were considered significant (i.e. HDI does not overlap 0).

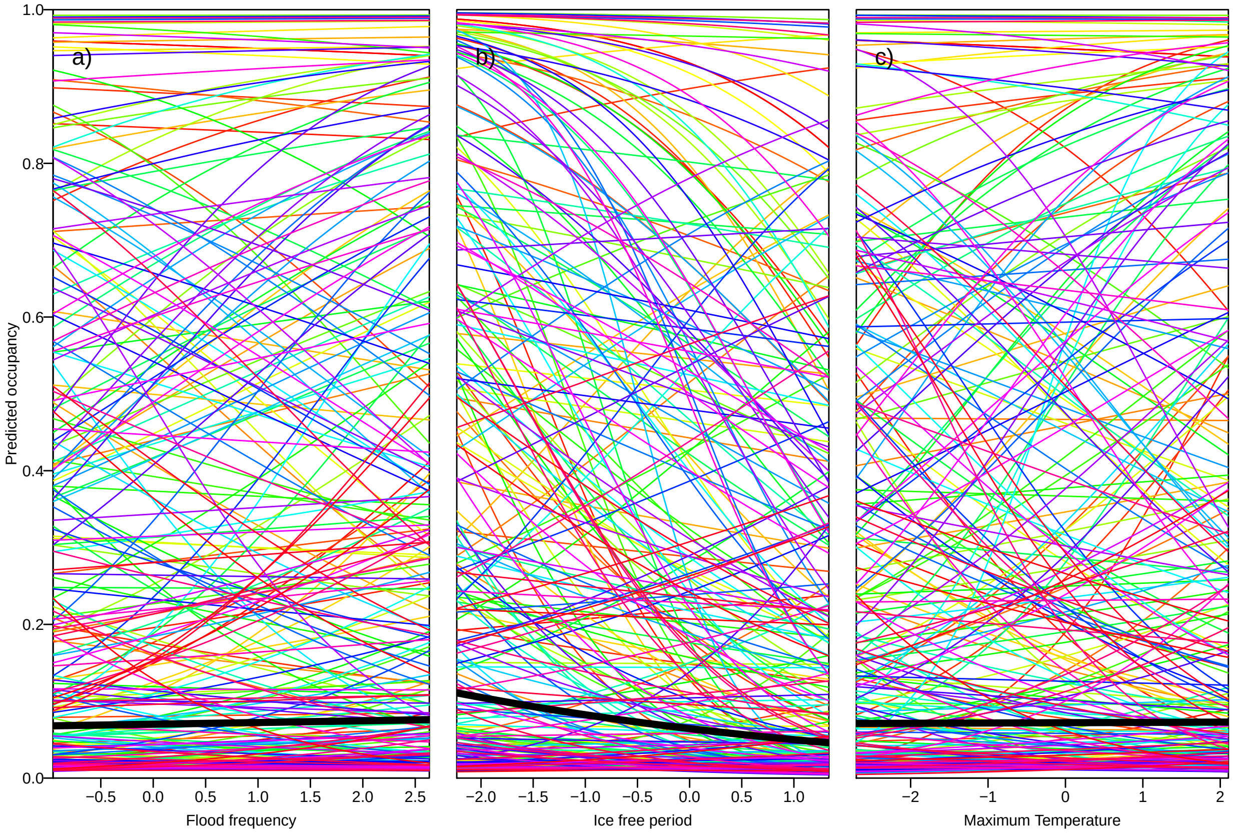


**Figure S1.13:** Impact on occupancy of specific genera (coloured lines) and the community on average (black line) due to changes in a) site flood frequency, b) the ice free period (prior to survey), and c) maximum temperature (prior to survey that year).

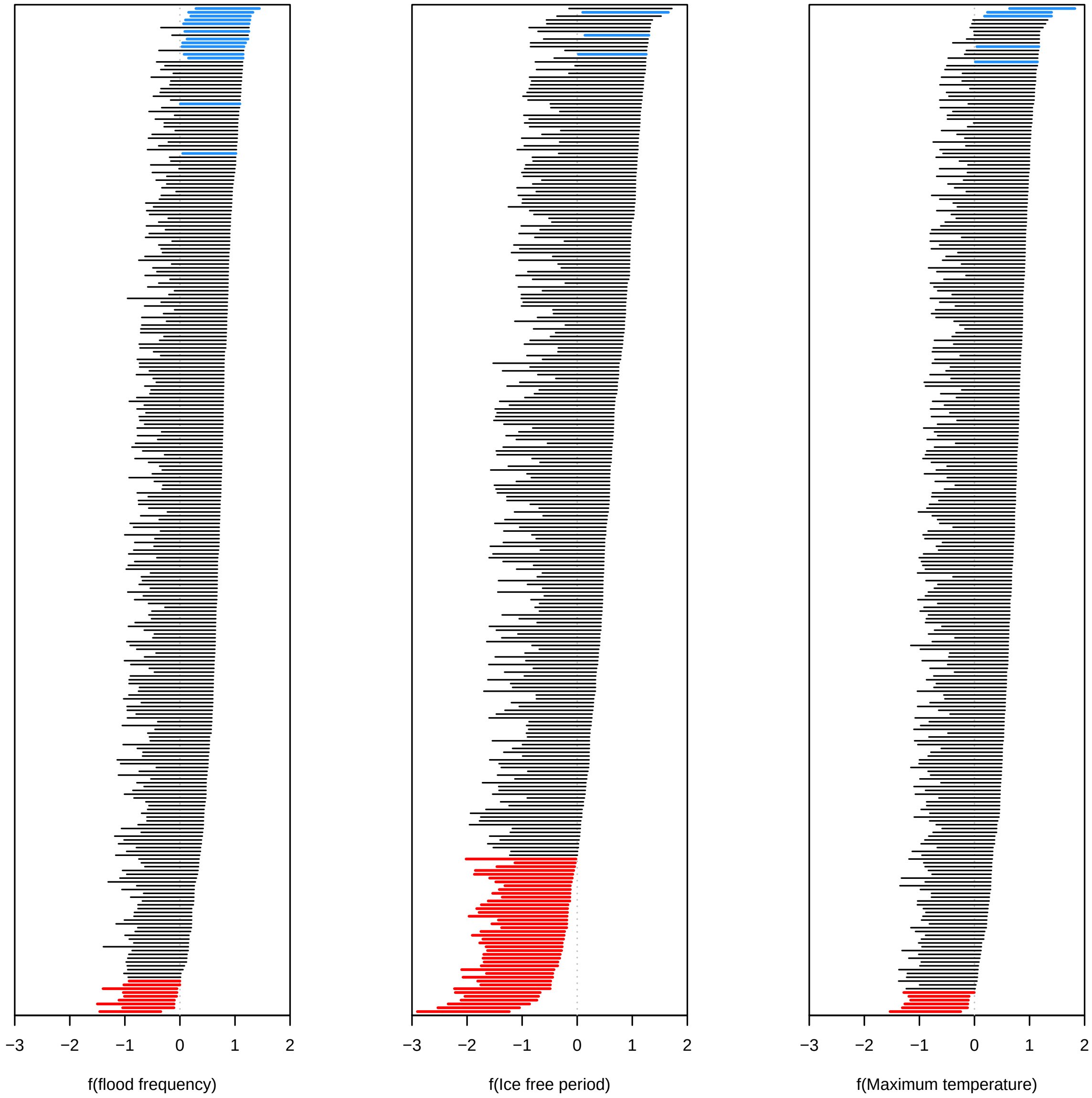


**Figure S1.14:** Highest density intervals (95%) of coefficients for the three environmental factors with significant positive (blue) and negative (red) effects on the occupancy of genera. Note each panel has been ordered for each coefficient separately and therefore the order of taxa varies in each case.

The genera that responded positively to the frequency of flooding were:

```
## [1] "Angarotipula"  "Banksiola"     "Cloeon"        "Dicrotendipes"
## [5] "Lecane"        "Polyphemus"    "Procladius"    "Triaenodes"
## [9] "Vejdovskyella"
```

The genera that responded negatively to the frequency of flooding were:

```
##  [1] "Acanthodiaptomus" "Carrhydrus"       "Gyrinus"
##  [4] "Laccophilus"      "Limnodrilus"      "Microvelia"
##  [7] "Motobdella"       "Neoporus"         "Phaenopsectra"
## [10] "Piona"            "Plumatella"       "Tubifex"
## [13] "Unionicola"
```

The genera that responded positively to the length of the ice free period were:

```
##  [1] "Aeshna"          "Agabus"          "Aglaodiaptomus" "Anabolia"
##  [5] "Callicorixa"     "Camptocercus"    "Carrhydrus"     "Cladopelma"
##  [9] "Coelambus"       "Corynoneura"     "Culex"          "Daphnia"
## [13] "Dytiscus"        "Eucypris"        "Eurycercus"     "Gerris"
## [17] "Glyptotendipes" "Graphoderus"     "Hesperocorixa"  "Hydaticus"
## [21] "Hydrobius"       "Hygrotus"        "Laccophilus"    "Ladislavella"
## [25] "Lejops"          "Lestes"          "Lumbriculus"    "Microvelia"
## [29] "Motobdella"      "Mytilina"        "Palpomyia"      "Piona"
## [33] "Plumatella"      "Polypedilum"     "Rhantus"        "Sigara"
## [37] "Simocephalus"    "Siphlonurus"     "Sympetrum"      "Tanysphyrus"
## [41] "Thulinius"
```

The genera that responded negatively to the length of the ice free period were:

```
## [1] "Ilybius"     "Nemotaulius" "Phryganea"
```

The genera that responded positively to the maximum temperature prior to survey were:

```
## [1] "Pristina"                    "Sida"
## [3] "Spongilla"                   "Stilobezzia"
## [5] "undef_Chironomidae"          "undef_undef_Sarcoptiformes"
```

The genera that responded negatively to the maximum temperature prior to survey were:

```
## [1] "Aglaodiaptomus" "Agraylea"       "Hedriodiscus"   "Ilybius"
## [5] "Ladislavella"
```

---

#### Compositional Turnover (beta diversity)

Given the effect detectability has on the expected 'true' number of species present at a site (alpha diversity) or in the metacommunity (gamma diversity), we would also expect detectability to affect our perception of compositional dissimilarity between sites (beta diversity). Dissimilarity may increase among sites if the majority of taxa that were not detected are also not prevalent, and hence unlikely to be shared among samples. Conversely, dissimilarity may decrease if the taxa that were not detected were in fact relatively common, and hence the probability that two sites do in fact share that taxon increases.

Figure S1.15 illustrates the differences between our original measures of dissimilarity, and the estimates we can now make from the four models based on each dataset, either for inter-annual dissimilarity at a site (left column), or among-site dissimilarity during a single visit (right column). This figure highlights a few key points:

1. Based on the CABIN dataset (a,b,e,f), imperfect detection was leading to an overestimation of pairwise dissimilarity among samples.
2. Based on DNA-metabarcoding (c,d,g,h), estimates from models are similar to the original values, particularly for among-site comparisons. This would suggest that even though large numbers of taxa were missed (see figures S8 & S9), the proportion that are shared or unique to a site remained comparable.
3. Compositional turnover increases as taxonomic resolution increases (c & g vs. d & h).

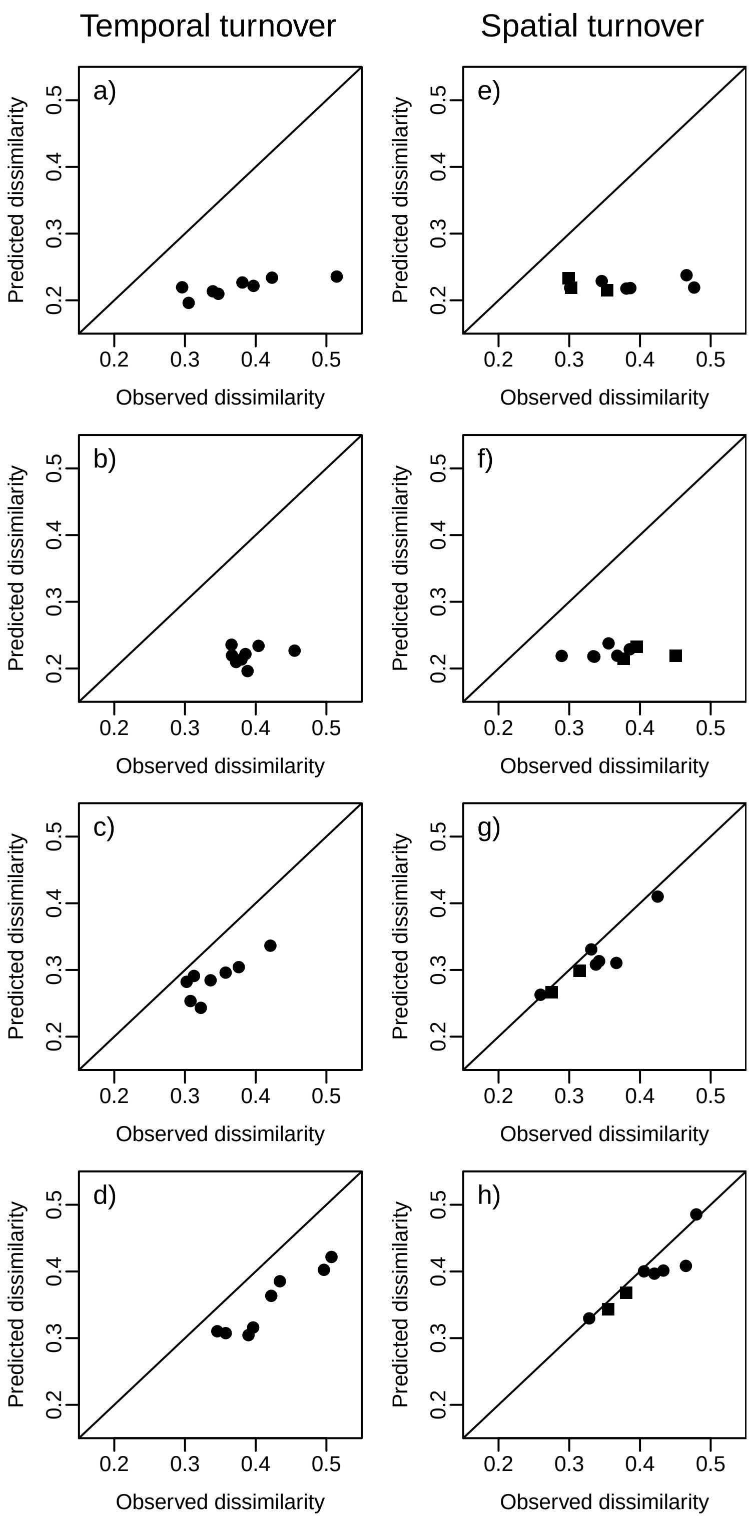


**Figure S1.15:** Comparison of the mean pairwise dissimilarity (Sorenson) observed in a community and predicted after accounting for detectability. The left column (a-d) compares dissimilarity among samples from the same site in different years, and the right column (e-h) the mean dissimilarity among sites during a single survey (year/season). Rows correspond to the four datasets analysed: CABIN presence-absence data (a,e), CABIN count data (b,f), and DNA metarcoded data at the family (c,g) and genus-level (d,h).
