## Supplement 2 for "Metabarcoding a Metacommunity: detecting change in a wetland wilderness"

### Supplementary Material S2: Metacommunity simulation and power analysis of macroinvertebrate biomonitoring in the Peace-Athabasca Delta based on morphological identification and DNA metabarcoding.

*Alex Bush*

*26th June 2019*

---

This supplementary file describes a simulation of our ability to sample and observe change in the aquatic macroinvertebrate metacommunity based on the estimates of diversity, site occupancy and detectability derived from the models described in Supplement S1.

### Power analysis

The primary aim of biological monitoring is to provide managers with an indication of ecosystem health, alert them to potential degradation of the system, and given most ecosystems are subject to multiple stressors, attribute the most likely cause (*Bonada et al. 2006 Annual Review of Entomology*). Therefore the principle question guiding this research program has been whether significant shifts in the composition of aquatic invertebrates can be detected in the Peace-Athabasca Delta (PAD). We used power analysis to understand how the inherent properties of the PAD ecosystem are likely to affect the quality of the data we can observe, and how reliably changes could be interpreted as statistically significant departures from expectations.

The data properties that influence our statistical power to discriminate between a baseline community and a degraded one are not only dependent on the study design (location and number of samples), but also on the mode of observation, namely the comparison between morphologically-identified CABIN samples and samples processed with DNA metabarcoding. Consequently the analysis below describes how we can understand the relative effect of both sampling effort and processing method on uncertainty in monitoring the aquatic invertebrates of the PAD.

It is worth emphasizing that we do not know the true parameters that describe the aquatic invertebrate metacommunity in the PAD ecosystem. However, by using the hierarchical models described in Supplement 1, we do have an estimate of the true *process model* and the *observation model* that governs what data we expect to recover. Therefore the power analysis investigates the sampling required to detect changes to the underlying process model, given our understanding of the observation processes.

To make comparisons based on sampling design and sample processing, we needed to keep the process-model consistent. As such we chose the model at genus-level to be the basis for simulated communities because DNA-metabarcoding encompassed most of the diversity observed by CABIN and genera could be aggregated to represent coarser taxonomic reoslution in the other models. Simulating the true distribution of abundance in invertebrate communities carried far greater uncertainty as the proportion of individuals sampled is unknown, and there was no straightforward way to compare occupancy and abundance. Hence the power analysis did not include the N-mixture model based on CABIN abundance data.

---

#### The metacommunity

Based on estimates of overall invertebrate genera-diversity (gamma) and occupancy derived from the multi-species occupancy model (MSOM), it is simple to reproduce a presence-absence matrix of the metacommunity from which we can take hypothetical samples. It can however be much more challenging to understand how diversity is distributed in space, and how quickly community composition will change over time. Although flood frequency does relate to geographic location, we regard it, and the other covariates related to temperature, as indicators of temporal change (Figure S2.1). Thus in any given generation of the simulation, occupancy of each taxon in the metacommunity is determined by the environmental conditions proposed (Figure S2.2). Whilst differences in occupancy do result in temporal turnover (approximately 10-27%), this is still not sufficient to explain the rate of turnover observed in the PAD. As a result, we simulated dynamic changes in local community composition (local extinction/colonization) that overlaid the shifts in occupancy due to environmental variation.

Turnover of the presence-absence matrix could be simulated using permutation of occurrences among sites in the simulated metacommunity. This could be continued until the rate of turnover approximated the degree of turnover we observed in the PAD data. For further information about the algorithms available to generate these changes, users should refer to the manual for the function *permat* in the vegan package. Note that although occupancy was allowed to vary between time steps, we preserved the row and column sums during permutation, meaning occupancy and site richness were held constant.

Ultimately, the genus-level turnover in the PAD was so high, on average above 40% (see Figure S1.15), that those observations could only be replicated by the complete redistribution of occurrences. Community assembly at an individual site could therefore be considered purely the result of chance events, with composition of any given site a product of its richness and the relative occupancy of taxa among sites.

Full permutation of presence-absence matrices to generate random assembly patterns is consistent with neutral assembly processes (Hubbell, 2001), although given the rate of turnover it is unlikely that dispersal is a limiting factor for the majority of taxa. The near-random dynamics of community assembly could instead be derived from a more complex pattern of neutral processes (random dispersal and stochastic local extinction), niche preferences (supported in part by the significant effects of environmental covariates on the occupancy of some taxa), patch dynamics and mass effects across the PAD landscape (Leibold & Chase 2018 *Metacommunity Ecology*).

To begin, the first piece of information to structure simulated PAD communities is their overall diversity (i.e. regional gamma diversity): `{r} M`

We then create a matrix to represent 100 possible wetland sites, and their environmental conditions over a number of timesteps, in this case representing years. A hundred sites was large enough to distribute species among sites according to their expected occupancy, but not so large as to be challenging to process. Likewise 20 timesteps was sufficient to test the power to observe any given change without storing and processing excessive volumes of data.

To generate hypothetical environmental conditions for this simulation we used the mean and covariance of the three variables in the original data to simulate new possible values. **Note** that these environmental conditions therefore reflect the data available for the 8 sites in this study, but may not be representative of the variation among all wetlands in the PAD, or of variation outside the six years for which we have data. The three variables that we included in the power analysis were flood frequency, time since ice melt and maximum temperature (see Supplement 1 for more details).

```r
library(jagsUI)
library(readxl)
library(knitr)
```

```r
library(betapart)
library(ecodist)
library(vegan)
library(mvtnorm)
library(mvabund)

# No. of sites
N            = 100
# No. of timesteps
TX           = 20
# Create a blank matrix to record starting occurrence
metacomm     = matrix( 0, nrow=N, ncol=M, dimnames=list(paste0("Site",c(1:N)), pad.lists$Species) )
# Standardize
standardize = function(x){ (x - mean(x))/sd(x)  }
Xcov         = apply(Xcov, 2, standardize)
Xcov         = rmvnorm(mean = apply(Xcov, 2, mean), sig = cov(Xcov), n=TX)
```

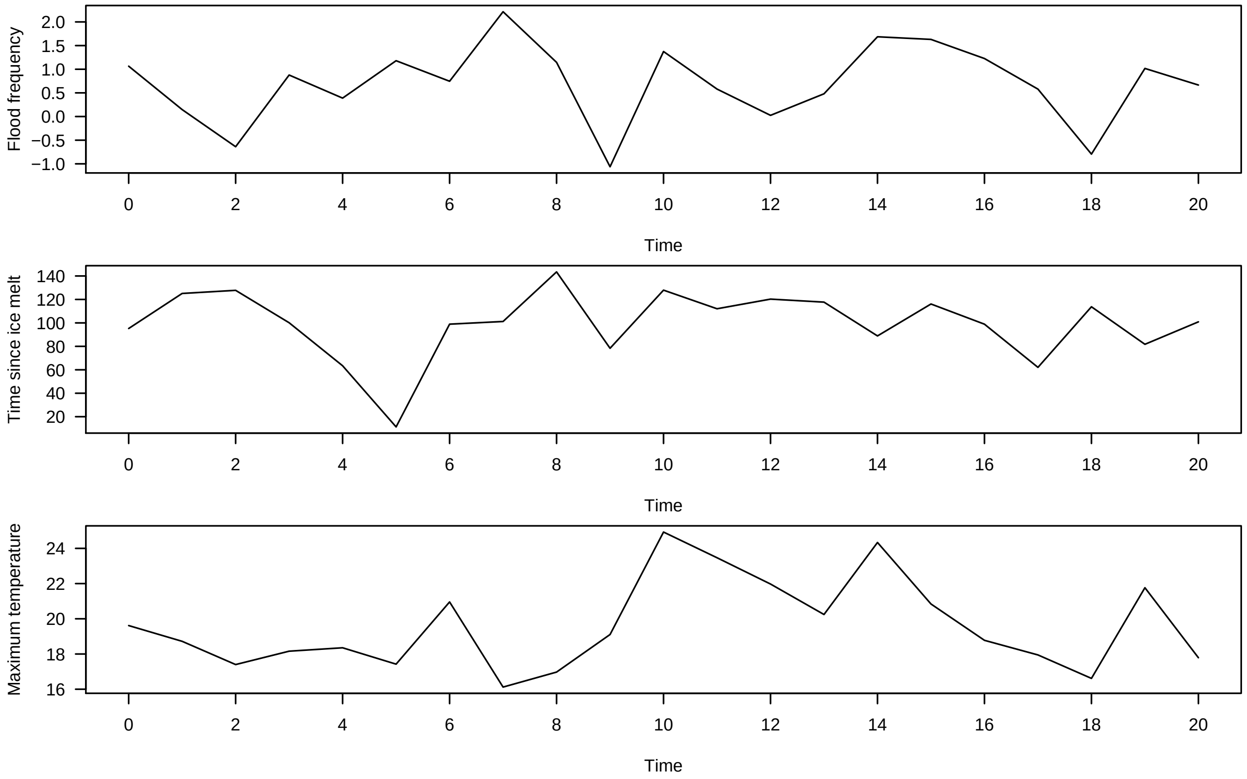

**Figure S2.1:** Example of a simulated timeseries of the three environmental covariates to determine the expected occupancy of taxa in the genus-level model (see Supplment 1 for more details). Covariates included flood frequency, the time since ice melt, and the maximum temperature prior to survey.

```r
# Create empty table for each species in each time step
spp.occ         = matrix(rep(0,M*TX), nrow=M, ncol=TX)
# Load fitted coefficients
beta            = cbind(MSOM$mean$lpsiS[1:M], MSOM$mean$betalpsiS[1:M,] )

# Fill-in expected occupancy (transforming predictions form logistic scale)
for(jj in 1:M){ spp.occ[jj,] = plogis(beta[jj,1] + (beta[jj,2]*Xcov[,1]) + (beta[jj,3]*Xcov[,2]) + (beta
```

```
# Plot
par(mfrow=c(5,5),oma=c(2,2,0.1,0.1),mar=c(2,2,0.5,0.5))
cols= rainbow(M)
xi = rep(seq(1,25,1),M)[1:M]
for(i in 1:25){
  plot(rep(0,TX) ~ c(1:TX), type="l", col="white", ylim=c(0,1),ylab="",las=1,xlab="")
  for(ii in which(xi==i)){ lines(spp.occ[ii,] ~ c(1:TX), col=cols[ii],lwd=2) }
}
mtext("Simulated generations (years)",1,line=0.5,outer=T)
mtext("Predicted occupancy",2,line=0.5,outer=T)
```

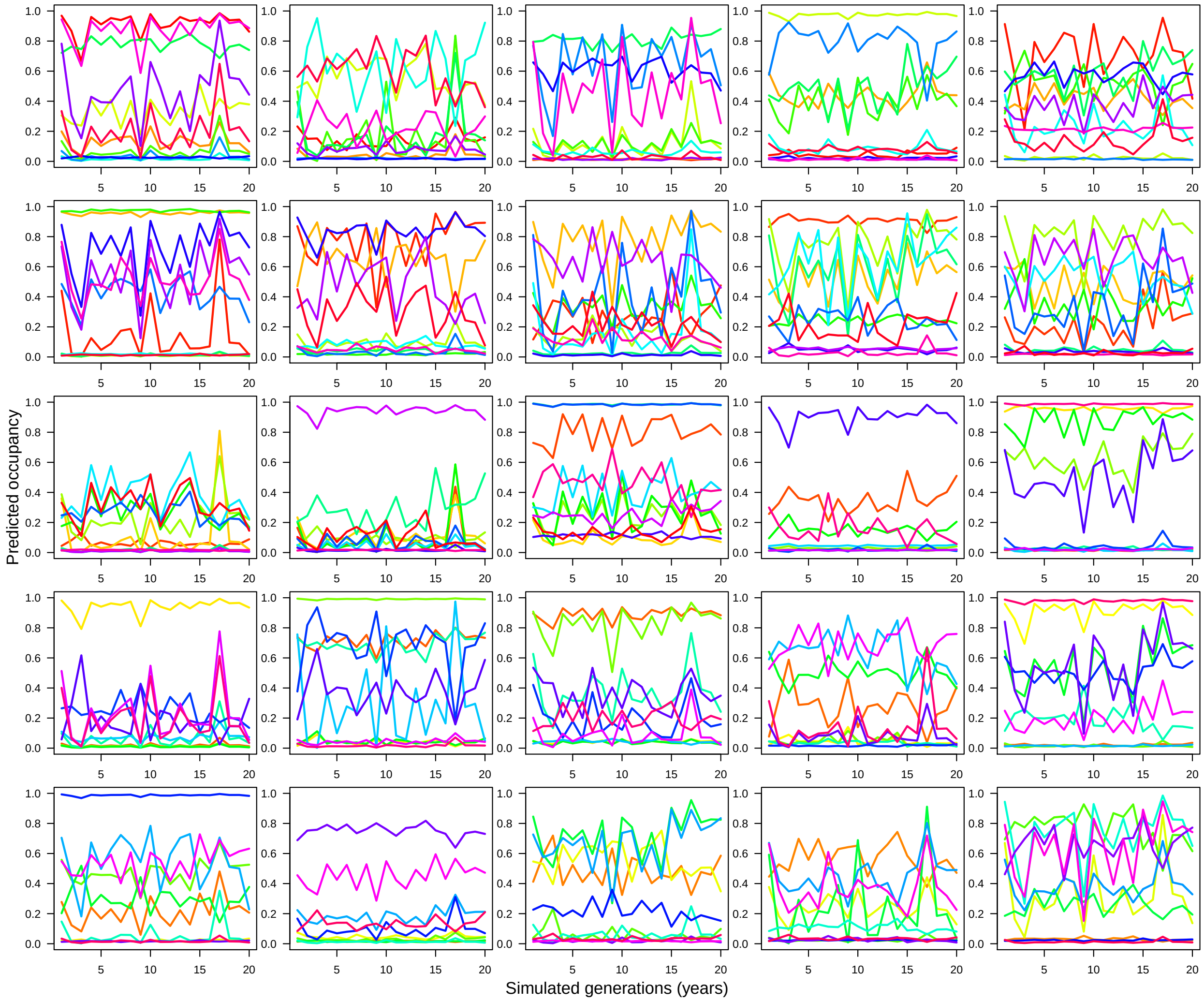

**Figure S2.2:** Simulated occupancy of 263 genera in response to changes in three environmental covariates over a twenty year period. Parameters were estimated by the hierarchical occupancy model described in Supplement 1, based on samples processed with DNA metabarcoding.

```
# Check all species always occur in at least one site, and if by chance they do not, increase so they d
spp.occ[round(spp.occ,2)==0] = 1/N
# Convert proportions to integers of occurrences
spp.occ = round(spp.occ,2) * N

# Create starting values to 'metacomm' for 1st timestep
```

```
for(i in 1:M){ metacomm[sample(c(1:N),spp.occ[i,1]),i] = 1 }

# And, working backwards, calculate the proportional change relative to previous generation
occ.delta = spp.occ
for(i in TX:2){ occ.delta[,i] = apply(occ.delta[,c(i-1,i)],1,diff) / (occ.delta[,i-1]/100) }
# There is no change in the first generation
occ.delta[,1] = 0
```

At this stage we reviewed the compositional dissimilarity among samples in space and time expected purely as a result of different probabilities of occurrence and the changes in those probabilities due to environmental responses (Figure S2.3). The turnover among sites in a simulated community (red solid line) falls within the distribution of dissimilarity values observed in the survey data (solid black line). However, the temporal turnover in the simulated community (dashed red line), driven solely by occupancy responses to fluctuating environmental conditions, was typically much lower than the temporal turnover observed in the PAD samples collected (dashed black line). We therefore introduced a process of permutation for the matrix of taxon occurrences which simulates temporal turnover through local extinction and colonisation, but maintains the known distribution of occupancy in the community.

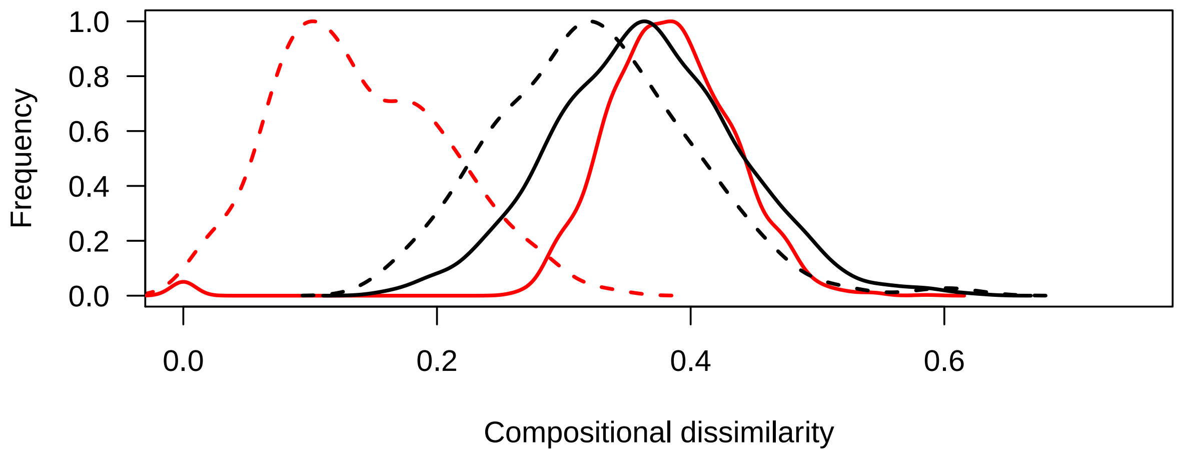

**Figure S2.3:** Distribution of pairwise dissimilairty among sites in space (solid lines) and over time (dashed lines) for the sites surveyed in the PAD (black lines), and the simulated community composition that replicates the same patterns of occupancy (red lines).

---

#### Sensitivity to Impact

As described in the main text, a challenge for estimating our ability to detect a deviation from the expected composition is that we do not know what form that deviation will take. A stressor is unlikely to drive the decline of a community at random. Furthermore, our study was meant to anticipate potential risks, and therefore we could not easily quantify the magnitude of effects that might be imposed on the ecosystem, such as the distribution and concentration of a given pollutant, or the expected changes in temperature and rainfall patterns. In addition, there is very little information available (e.g. trait data, ecotoxicological trials, field experiments) for how the taxa in this system might respond, particularly for diverse assemblages like invertebrates.

Given we could not use empirical data to assign realistic thresholds, or even approximate the parameters that govern the sensitivity of specific taxa, we modelled the degradation of the metacommunity in response to a hypothetical stressor, hereafter referred to as 'Stressor-X'. Values of stressor-X are thus meaningless. However, what is important is how sensitivity co-varies with taxon's occupancy. One extreme scenario would be for common taxa to also be the most tolerant, and hence identifying the impact of a stressor would rely on observing the loss of taxa with low occupancy (loss of rare taxa). Conversely, the opposite extreme proposed that the most prevalent taxa are the most sensitive to Stressor-X, and therefore the impact of stressor-X is related to the decline of common taxa.

What is missing from the description above, and most other studies, is a consideration of detectability. If species with low tolerance also have a low probability of detection, then changes to the community will be more difficult to confirm with confidence. However, as this study has already modelled the relationship between occupancy and detectability, these characteristics of the community are effectively known. Our statistical power to detect change therefore depends on the magnitude of a shift in composition (the "signal") relative to the background variability of the system (see discussion on fluctuating occupancy and permutation above) and sampling error ("noise" due to imperfect detection).

---

#### The simulation function

The simulation function took the following broad steps:
1. To begin, a metacommunity matrix, generated as described above using the parameters estimated by the hierarchical occupancy models, is provided to the function (*start.comm*). As this represents the expected metacommunity at its natural baseline equilibrium, it is referred to as the "null" model.
2. The overall occupancy of each taxon is adjusted in each of *TX* timesteps based on the proportional change described by the *occ.delta* arguement. After this, permutation of the occurrence matrix is used to randomize the assemblage structure and then stored in an array. The final array has *TX* layers equivalent to *start.comm* matrix. This array represents the *null* metacommunity.
3. Repeat the process for a copy of the metacommunity matrix, and then further remove occurrences of taxa whose assigned threshold tolerance (see *SSDx*) falls below the value of Stressor-X (specified by *xgradient*). This array represents the *disturbed* metacommunity.
4. If an argument for *higher.taxa* is provided, the arrays for *null* and *disturbed* occurrence are aggregated to reflect occurrence of their parent taxonomic rankings; in this case genus to family-level.
5. New matrices are then created to represent the **observed** composition of both the *null* and *disturbed* metacommunities given a) the detectability of the taxa, and b) the number of samples taken.
6. Finally the compositional dissimilarity of the surveyed sites in the *null* and *disturbed* arrays is compared in each annual time step using *mvabund*. For the purposes of this study we were interested in how many samples would be required to detect a particular ($p$<0.05) shift in composition with a given degree of confidence. The number of samples required to detect shifts in community composition in any given year was very high. As a result, for the purposes of this study we studied the power required to detect a significant shift at least 50% of the time (i.e. on average within 2 years). Significant differences therefore needed to be observed in half the TX steps to be considered successful. As this testing phase was the most time consuming step, a further argument (*mva.stop*) was added to the routine so that the function would escape if this desired threshold was met (i.e. the no. of significant results was equivalent to TX/2).

```
metacomm.permute = function(TX,                  # Total no. of time steps (years)
                            start.comm,          # Starting compositional matrix
                            occ.delta,           # Proportional change (+/-) in occupancy.
                            higher.taxa = NULL, # Table of taxon names and higher ranks.
                            # Disturbance parameters
                            SSDx      = NULL,    # Species Sensitivity Distribution
                            xgradient = NULL,    # Vector of 'StressorX'.
                            # Observation parameters
                            survey,              # Index of surveyed sites (rows)
```

```
                           detection = NULL,   # List of detectabilities and reps
                           mva.stop  = NULL
){
  # No. of taxa and sites
  M = ncol(start.comm)
  N = nrow(start.comm)

  # - - - - - - - - - - - - - - - - - - - - - - - - - - - - #
  # CREATE NULL METACOMMUNITY
  # - - - - - - - - - - - - - - - - - - - - - - - - - - - - #

  # Create null array to record composition (site X species) in TX annual time steps
  tcomp = array(0, dim=c(N,M,TX), dimnames=list(paste0("Site",c(1:N)),
                                                paste0("Spp", c(1:M)),
                                                paste0("Time",c(1:TX))) )
  # Composition at time 1 is equivalent to 'start.comm'
  tcomp[,,1] = start.comm

  # Loop over time steps
  for(i in 2:TX){
    # Start with composition of previous time-step
    comp = tcomp[,,i-1]
    # Account for any changes in species' occupancy
    for(ii in which(occ.delta[,i]!=0) ){
      # Note: proportional changes must be rounded to nearest whole number
      xi = round((sum(comp[,ii])/100) * (occ.delta[ii,i]))
      if( xi>0 ){
        xii = which(comp[,ii]==0)
        xi  = min(xi,length(xii))
        comp[sample(xii,abs(xi)),ii] = 1
      }
      if( xi<0 ){
        xii = which(comp[,ii]==1)
        xi  = min(abs(xi),length(xii))
        comp[sample(xii,abs(xi)),ii] = 0
      }
    }
    # Permutation
    X = permatswap(comp,method="tswap",fixedmar="both",mtype="prab",
                   times=1,burnin=150000,thin=1000)
    # Enter matrix with new composition into the next timestep of array
    tcomp[,,i] = X$perm[[1]]
  } # End TX loop

  # - - - - - - - - - - - - - - - - - - - - - - - - - - - - #
  # OVERLAY DISTURBANCE ON EQUIVALENT METACOMMUNITY
  # - - - - - - - - - - - - - - - - - - - - - - - - - - - - #

  # Create an equivalent metacommunity array to record responses to Stressor-X.
  dcomp     = tcomp
  dcomp[,,] = 0
  # If stress isn't specified for each year, use the first value for all years
  if(length(xgradient)!=TX){
```

```
  xgradient = rep(xgradient[1],TX)
  print("Using first value of xgradient for all TX years.")
}

# Loop over time steps
for(i in 1:TX){
  # Start with composition of previous time-step
  # Note unlike the null routine above is that we now loop over step 1 too
  # and hence must make sure here that we don't refer to a previous timestep.
  if(i>1){ comp = dcomp[,,i-1] } else { comp = start.comm }

  # Account for any changes in species' occupancy
  for(ii in which(occ.delta[,i]!=0) ){
    # Note: proportional changes must be rounded to nearest whole number
    xi = round((sum(comp[,ii])/100) * (occ.delta[ii,i]))
    if( xi>0 ){
      xii = which(comp[,ii]==0)
      xi  = min(xi,length(xii))
      comp[sample(xii,abs(xi)),ii] = 1
    }
    if( xi<0 ){
      xii = which(comp[,ii]==1)
      xi  = min(abs(xi),length(xii))
      comp[sample(xii,abs(xi)),ii] = 0
    }
  }

  # Remove occurrences from community below a certain threshold
  # Which taxa are below the threshold for year 'i'?
  xid = which(SSDx < xgradient[i])
  # Set occurrences to zero
  comp[,xid] = 0
  # Permutation
  X = permatswap(comp,method="tswap",fixedmar="both",mtype="prab",
                 times=1,burnin=150000,thin=1000)
  # Store output
  dcomp[,,i] = X$perm[[length(X$perm)]]

} # End TX loop

# - - - - - - - - - - - - - - - - - - - - - - - - - - - #
# AGGREGATE THE GENUS-LEVEL COMMUNITY TO FAMILY-LEVEL TAXA
# - - - - - - - - - - - - - - - - - - - - - - - - - - - #

if( !is.null(higher.taxa) ){
  # 'HR' is for 'higher rank'. Create arrays the right size
  HRtcomp     = tcomp[,1:length(unique(higher.taxa[,1])),]
  HRdcomp     = dcomp[,1:length(unique(higher.taxa[,1])),]

  # Now aggregate occurrences to higher taxon rank groups.
  for(i in 1:TX){
    HRtcomp[,,i] = t(aggregate(t(tcomp[,,i]), by=list(higher.taxa[,1]), FUN=max)[,-1])
```

```
    HRdcomp[,,i] = t(aggregate(t(dcomp[,,i]), by=list(higher.taxa[,1]), FUN=max)[,-1])
  }
  tcomp = HRtcomp ; rm(HRtcomp)
  dcomp = HRdcomp ; rm(HRdcomp)
}

# - - - - - - - - - - - - - - - - - - - - - - - - - - - #
# APPLY DETECTION ERROR TO EACH METACOMMUNITY
# - - - - - - - - - - - - - - - - - - - - - - - - - - - #

# Arguement 'detection' includes detectability with each mode of observation and the
# number of replicate samples using that method.
det1  = detection[[1]]
reps1 = detection[[2]]
det2  = detection[[3]]
reps2 = detection[[4]]

# Overlap detectabilities on occurrence matrix based on different levels of
# sampling ('reps') and processing effort (separate primers)
tcomp.obs = tcomp
dcomp.obs = dcomp
for(i in 1:TX){
  if( reps1>0 ){
    simcomm.t1 = rbinom(tcomp[,,i], reps1, det1) * tcomp[,,i]
    simcomm.d1 = rbinom(dcomp[,,i], reps1, det1) * dcomp[,,i]
  } else {
    simcomm.t1 = simcomm.d1 = matrix(0,N,M)
  }
  if( reps2>0 ){
    simcomm.t2 = rbinom(tcomp[,,i], reps2, det2) * tcomp[,,i]
    simcomm.d2 = rbinom(dcomp[,,i], reps2, det2) * dcomp[,,i]
  } else {
    simcomm.t2 = simcomm.d2 = matrix(0,N,M)
  }
  # Combine
  simcomm.t = simcomm.t1 + simcomm.t2 ; simcomm.t[simcomm.t>0] = 1
  simcomm.d = simcomm.d1 + simcomm.d2 ; simcomm.d[simcomm.d>0] = 1
  # Fill array
  tcomp.obs[,,i] = simcomm.t
  dcomp.obs[,,i] = simcomm.d

  rm(simcomm.t1,simcomm.t2,simcomm.d1,simcomm.d2,simcomm.t,simcomm.d)
}

# - - - - - - - - - - - - - - - - - - - - - - - - - - - - - #
# SUMMARY STATS
# - - - - - - - - - - - - - - - - - - - - - - - - - - - - - #

# Test for differences between null and degraded communities with mvabund

# Can compare groups of sites from A SINGLE year:
mv.anova = data.frame("TX"=1:TX,"Dev.null"=NA,"p.null"=NA,"Beta.null"=NA,
                      "Dev.obs"=NA,"p.obs"=NA,"Beta.obs"=NA)
```

```
  if(is.null(mva.stop)){ mva.stop = TX }

  for(i in 1:TX){
    # The 'mva.stop' argument specifies an escape so that the function stops calculating significance a
    if( sum(mv.anova$p.obs[!is.na(mv.anova$p.obs)]<=0.05)<mva.stop ){
      inv.spp = mvabund(rbind(tcomp[survey,,i],dcomp[survey,,i])) # Format data for 'mvabund'
      xgroup  = factor(rep(c("A","B"),each=length(survey)))       # Create 'treatement' factor
      xi      = manyglm( inv.spp ~ xgroup, family="binomial")
      xii     = anova(xi)
      rowindex= c(1:length(survey))+length(survey)
      nrow    = length(survey)*2
      colindex= c(1:length(survey))
      Beta    = mean( as.matrix(vegdist(inv.spp))[rowindex + nrow * (colindex-1)])
      mv.anova[i,c("Dev.null","p.null","Beta.null")] = c(xii$table$Dev[2],xii$table$`Pr(>Dev)`[2],Beta)

      # Repeat for observed matrices
      inv.spp = mvabund(rbind(tcomp.obs[survey,,i],dcomp.obs[survey,,i]))
      xi = manyglm( inv.spp ~ xgroup, family="binomial")
      xii = anova(xi)
      Beta = mean( as.matrix(vegdist(inv.spp))[rowindex + nrow * (colindex-1)])
      mv.anova[i,c("Dev.obs","p.obs","Beta.obs")] = c(xii$table$Dev[2],xii$table$`Pr(>Dev)`[2],Beta)
    }
  }

  # - - - - - - - - - - - - - - - - - - - - - - - - - - - - - -  #
  # SAVE
  # - - - - - - - - - - - - - - - - - - - - - - - - - - - - - -  #

  # Save as list
  tdat = list("TX"          = TX,
              "N"           = N,
              "M"           = M,
              "Stress"      = list("SSDx"      = SSDx,
                                   "xgradient" = xgradient),
              "Survey"      = survey,
              "MVA"         = mv.anova,
              "Permutation" = list(tcomp,tcomp.obs,dcomp,dcomp.obs) )

  # - - - - - - - - - - - - - - - - - - - - - - - - - - - - - -  #
  # - - - - - - - - - - - - - - - - - - - - - - - - - - - - - -  #

  return(tdat)
}
```

---

For the purposes of this example we wished to illustrate a case where the degree of degradation in the metacommunity was not consistently detected in every single year based on the same sample size (8 sites) as was available in our dataset. We therefore ran the code below applying a stressor threshold that reduced the overall number of occurrences in the metacommunity by 18% (see *Stress.prop*=0.18), and specify a neutral relationship (*SSD.cor*=0) between occupancy and tolerance to Stressor-X.

```
# - - - - - - - - - - - - - - - - #
# SAMPLE SIZE
# - - - - - - - - - - - - - - - - #
```

```r
# As example, sample 8 of the N sites (rows) in metacomm
Survey.sites = sample(c(1:N),8)

# - - - - - - - - - - - - - - - - #
# GET PREDICTED DETECTABILITY
# - - - - - - - - - - - - - - - - #

# Draw estimates of detectability from the occupancy model
det1      = matrix(rep(plogis(MSOM$mean$lpS[1:M,1]),e=N),nrow=N) # BR5
det2      = matrix(rep(plogis(MSOM$mean$lpS[1:M,2]),e=N),nrow=N) # F230

# Note that a detection argument of 'list(det1,1,det2,1)' indicates the community was sampled with each
# For comparison with morphological approaches we simply specify no samples in the last entry: 'list(de

# - - - - - - - - - - - - - - - - #
# SSD
# - - - - - - - - - - - - - - - - #

SSD.cor = 0

standard01.complement = function(y, rho, x) {
  range01 = function(x){ (x-min(x))/(max(x)-min(x)) }
  if(missing(x)){ x = rnorm(length(y)) } # Optional: supply a default if `x` is not given
  yp  = residuals(lm(x ~ y))
  q   = rho * sd(yp) * y + yp * sd(y) * sqrt(1 - rho^2)
  q   = range01(q)
  return(q)
}

SSDx = standard01.complement(y=spp.occ[,1], rho=SSD.cor, x=runif(M,0,1)) # SSDx vector with desired cor

# Identify the value of StressorX that affects the target proportion of the occurrences for the communi
# As the frqeuency of different tolerances varies the quantile is weighted by occupancy.
Stress.prop = 0.18
StressX     = unlist(lapply(SSDx, FUN=function(x) rep(x,e=as.numeric(colSums(metacomm))[match(x,SSDx)])
StressX     = quantile(StressX, Stress.prop)

# Run simulation

MP = metacomm.permute(TX,
                      start.comm    = metacomm,
                      occ.delta,
                      higher.taxa   = NULL,
                      SSDx,
                      xgradient     = rep(StressX,TX),
                      survey        = Survey.sites,
                      detection     = list(det1,1,det2,1),
                      mva.stop      = NULL)
```

```
## Time elapsed: 0 hr 0 min 16 sec
## Time elapsed: 0 hr 0 min 17 sec
```

```
## Time elapsed: 0 hr 0 min 16 sec
## Time elapsed: 0 hr 0 min 17 sec
## Time elapsed: 0 hr 0 min 16 sec
## Time elapsed: 0 hr 0 min 17 sec
## Time elapsed: 0 hr 0 min 16 sec
## Time elapsed: 0 hr 0 min 17 sec
## Time elapsed: 0 hr 0 min 16 sec
## Time elapsed: 0 hr 0 min 18 sec
## Time elapsed: 0 hr 0 min 18 sec
## Time elapsed: 0 hr 0 min 18 sec
## Time elapsed: 0 hr 0 min 18 sec
## Time elapsed: 0 hr 0 min 17 sec
## Time elapsed: 0 hr 0 min 17 sec
## Time elapsed: 0 hr 0 min 19 sec
## Time elapsed: 0 hr 0 min 18 sec
## Time elapsed: 0 hr 0 min 17 sec
## Time elapsed: 0 hr 0 min 16 sec
## Time elapsed: 0 hr 0 min 17 sec
## Time elapsed: 0 hr 0 min 16 sec
## Time elapsed: 0 hr 0 min 17 sec
## Time elapsed: 0 hr 0 min 17 sec
## Time elapsed: 0 hr 0 min 18 sec
## Time elapsed: 0 hr 0 min 16 sec
## Time elapsed: 0 hr 0 min 17 sec
## Time elapsed: 0 hr 0 min 16 sec
## Time elapsed: 0 hr 0 min 18 sec
## Time elapsed: 0 hr 0 min 17 sec
## Time elapsed: 0 hr 0 min 17 sec
## Time elapsed: 0 hr 0 min 16 sec
## Time elapsed: 0 hr 0 min 17 sec
## Time elapsed: 0 hr 0 min 16 sec
## Time elapsed: 0 hr 0 min 18 sec
## Time elapsed: 0 hr 0 min 16 sec
## Time elapsed: 0 hr 0 min 18 sec
## Time elapsed: 0 hr 0 min 17 sec
## Time elapsed: 0 hr 0 min 17 sec
## Time elapsed: 0 hr 0 min 17 sec
## Time elapsed: 0 hr 0 min 17 sec
```

---

#### Simulation results

To begin looking at how the outputs of the simulation were interpreted, Figure S2.4 summarises how the overall number of occurrences in the sampled sites were expected to vary naturally over a 20 year period. As this is the true state expected by the baseline surveys, we refered to it as the *Null process* model. The *Null observed* model shows the number of occurrences we would then expect to detect given the detectability of the taxa. Based on the simulated introduction of stressor-X, and an approximate 18% reduction in occurrences (see above), the red lines display the equivalent relative number of occurrences for the *Stressed process* model and *Stressed observed* model.

The red dots in Figure S2.4 indicate when there is a significant difference between the process data models, or observed data models, as identified by the multivariate generalized linear models (see *mvabund* package). As shown, with perfect detection and recovery of all taxa present, the differences in composition between the *null-process* and *stressed-process* models were consistently detected. Once we account for detectability the differences between the communities were captured more intermittently. As mentioned, this power analysis was used to identify at what value of *Stress.prop* were the differences in the observed data likely to be detected at least 50% of the time.

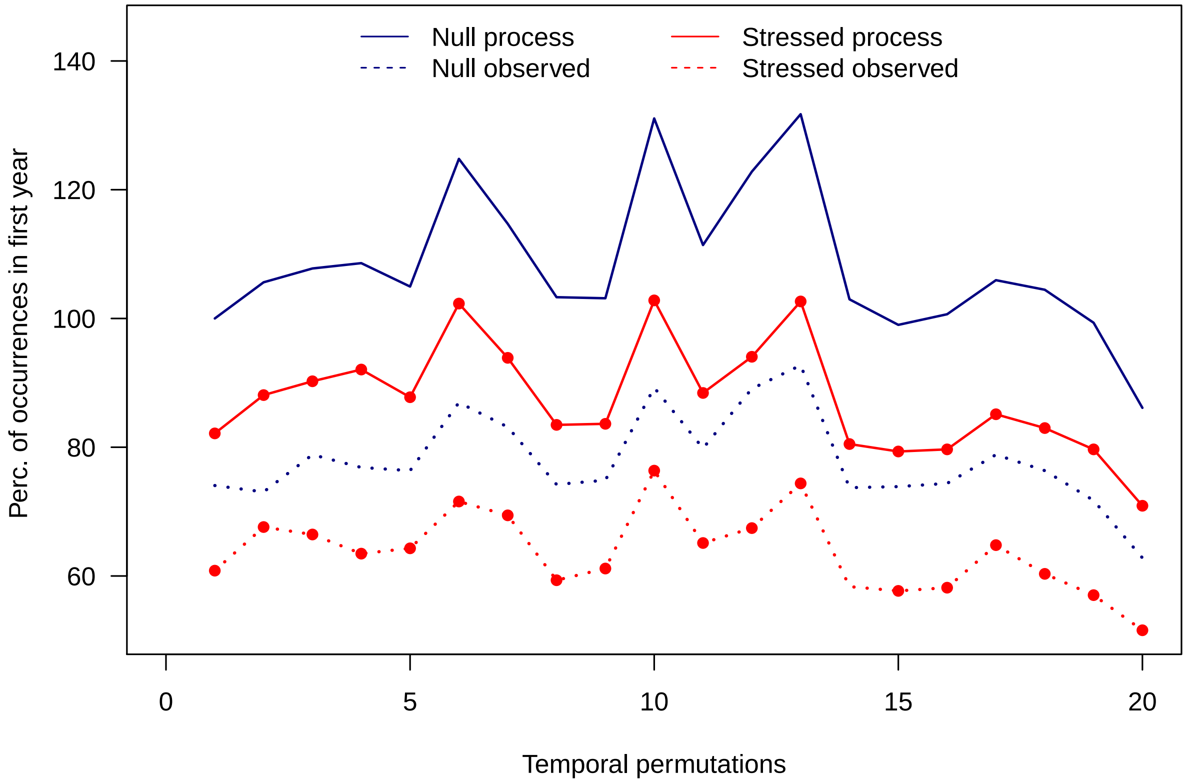

**Figure S2.4:** Proportion of occurrences sampled relative to the number present at the start of the simulated null community. The blue solid line represents the null model of the baseline metacommunity processes, and the red solid line depicts the same community impacted by a selected degree of stress, which in this case reduces the number of occurrences by 18%. The dotted lines represent the fraction of occurrences observed in samples from null and stressed communities. A red dot marks when the composition of the process or observed models were significantly different from each other.

---

Another option would be to compare how dissimilar the composition of *null* and *degraded* communities would be, and how dissimilar they appear to be after accounting for detectability.

```
XYLIM = range(MP$MVA$Beta.obs,MP$MVA$Beta.null)
plot(MP$MVA$Beta.obs ~ MP$MVA$Beta.null,xlim=XYLIM,ylim=XYLIM,
     xlab = "True dissimilarity of sampled sites",
     ylab = "Observed dissimilarity of sampled sites", las=1)
abline(0,1)
xi = which(MP$MVA$p.null<=0.05)
for(i in xi){ lines(x=rep(MP$MVA$Beta.null[i],2), y=c(MP$MVA$Beta.obs[i]-0.001,MP$MVA$Beta.obs[i]-0.005
xi = which(MP$MVA$p.obs<=0.05)
for(i in xi){ lines(x=c(MP$MVA$Beta.null[i]-0.001,MP$MVA$Beta.null[i]-0.005),
                    y=rep(MP$MVA$Beta.obs[i],2),lwd=2,col="red" ) }
```

```r
points(MP$MVA$Beta.obs ~ MP$MVA$Beta.null, pch=16)
```

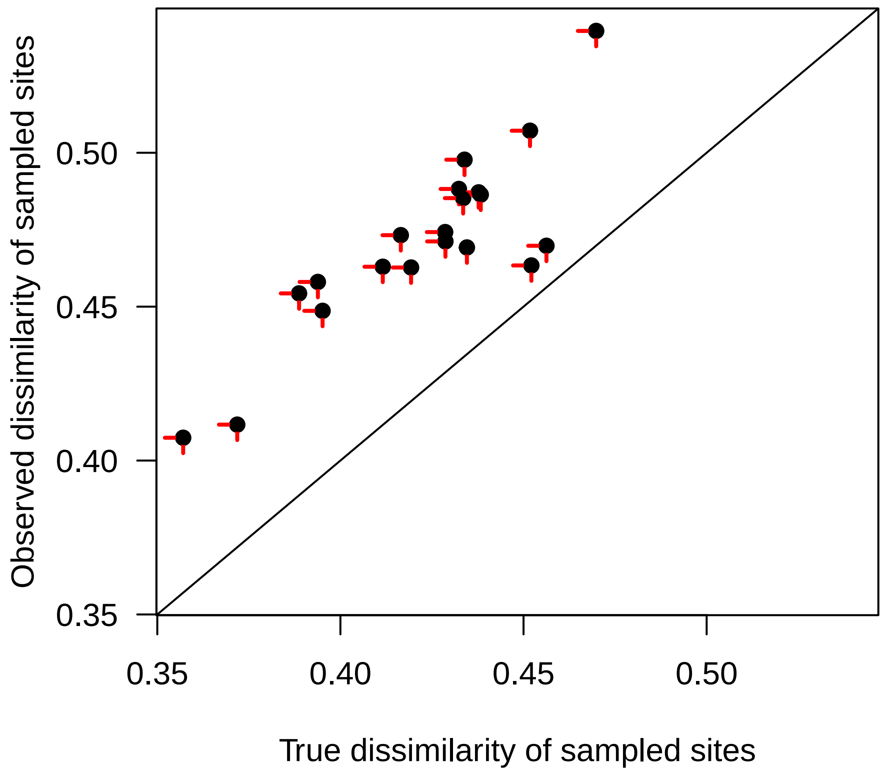

**Figure S2.5:** Relationship between the true and observed compositional dissimilarity between sampled sites for each simulated year. If composition was significantly different between the expected null community and the stressed equivalent a vertical red tick is added to the point (see points on solid red line in Figure S2.4). Likewise, significant differences between the metacommunities after accounting for detectability are indicated with a horizontal red tick (see points on dotted red line in Figure S2.4).

Given the observed spatial and temporal turnover in composition, it would be unrealistic to expect surveys to reliably detect differences from the expected null community in **any** year. Therefore, sampling was considered adequate if the stressed community was detectably different from the expected null community at least 50% of the time. We subsequently repeated the simulation 100 times for each combination of sample size and correlation with occupancy to account for variation in the assigned occupancy, tolerance thresholds and environmental parameters. If too many simulations failed to meet the desired threshold, the magnitude of Stressor-X was increased, further reducing the number of occurrences in the stressed community. The reduction in taxon occupancy was stopped once significant differences could be identified in 50% of years with 95% confidence, (i.e. in >95 simulations).

Below, Figure S2.6 describes how the threshold for detection was moderated by both sample size and the correlation with detectability. For example, a reduction in occurrences of approximately 10% could be detected by 7 samples per year **IF** a taxon's tolerance to a particular factor ('Stressor-X') was perfectly negatively correlated with its occupancy (i.e. common taxa are least tolerant, and hence first to be affected by increased stress). Conversely if rare taxa are most sensitive to stress (positive correlation in Figure S2.6 x-axis), then 12-14 samples may be required to detect the same 10% reduction in occupancy. Note that these sample sizes are not generic recommendations; they are specific to the PAD metacommunity, because the distribution of occupancy and correlation of occupancy with detectability will influence statistical power.

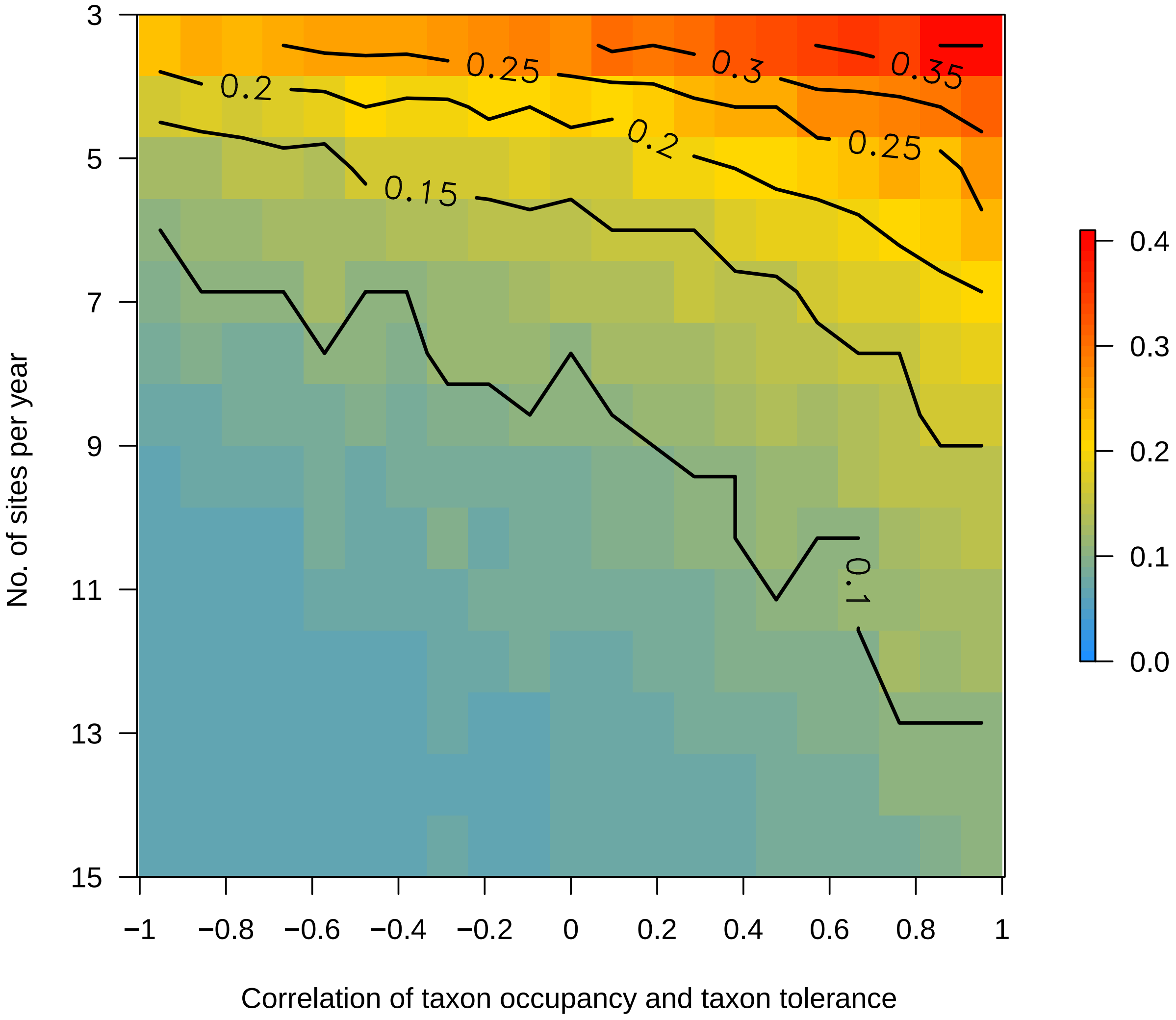

**Figure S2.6:** Minimum reduction to community occupancy detectable in 50% of years, with 95% confidence. Plot shows relationship between the number of sites sampled per year (a single sample per site), and the correlation of community tolerance with occupancy (see *Simulation to Impact* section above), upon the magnitude of change to the PAD metacommunity that can be detected.

Finally we compared the power to detect a reduction in occupancy, relative to the 'null' baseline metacommunity model, using different approaches to data generation. Figure S2.7 demonstrates that our power to confidently detect changes in occupancy increases both as sample size increases (either by increasing the number of sites per year or increasing within-site replication), and as the data resolution increases, specifically improving from CABIN family data to DNA-based family data, and again to the DNA data at genus-level. For example, even surveying 15-20 sites per year (a higher level of sampling effort than has been possible to date), we would not expect CABIN data to reliably detect reductions in occupancy less than about 20%, whereas DNA metabarcoding at the genus-level could detect reductions close to 5%.

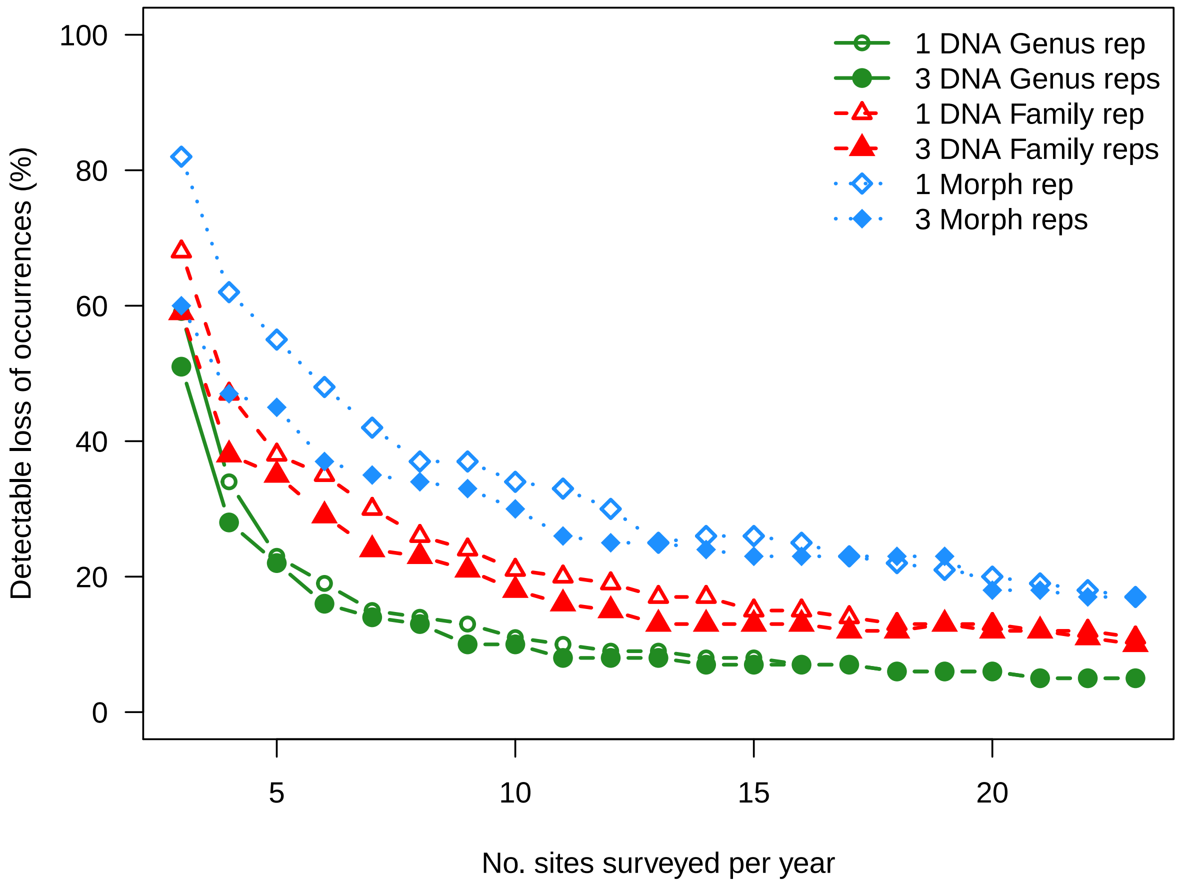

```
## pdf
##   2
```

**Figure S2.7:** Minimum detectable reduction of community occupancy (in 50% of years with 95% confidence) in response to number of annual survey sites. The threshold at which changes could be detected was influenced by approach to processing samples, the taxonomic resolution, and the number of replicate samples at each site surveyed.

---
